# The last one in the row but decisive: PPDK as a potential key regulator of diurnal deacidification in CAM leaves across varying PPFD and photoperiod conditions

**DOI:** 10.1101/2025.03.20.644348

**Authors:** Stijn Daems, Bram Van de Poel, Johan Ceusters

**Author notes:** Correspondence: Johan Ceusters.

## Abstract

- Crassulacean acid metabolism (CAM) plants primarily fix atmospheric CO_2_ at night and store it as malic acid in their vacuoles. During the light period, vacuolar malate is remobilised and decarboxylated to supply CO_2_ for Rubisco assimilation. Light intensity and photoperiod are believed to play crucial roles in regulating this process, but their influences on the underlying molecular and biochemical mechanisms remain unclear.
- In this study, we integrated physiological, biochemical, and molecular approaches to uncover the temporal patterns and light responsiveness of gene transcript and protein abundances, and the activities of enzymes involved in diurnal malate remobilisation in the obligate CAM model species *Kalanchoë fedtschenkoi*.
- Vacuolar malate transport was primarily influenced by the endogenous clock and photoperiod, with *Kf*ALMT4 being a more plausible transporter candidate than *Kf*tDT. Decarboxylation of the released malate was mainly dictated by photoperiod, with light intensity playing a supplementary role. Both photoperiod and light intensity greatly affected the final processes of CAM photosynthesis i.e. CO_2_ refixation and pyruvate recycling, with PPDK—the last in line—being the most strictly light-regulated player at the mRNA, protein abundance and activity levels, closely matching malate dynamics.
- Collectively, this study revealed the recycling enzyme PPDK as a potential key regulator of light-dependent diurnal deacidification in CAM leaves, rather than the vacuolar malate transport or decarboxylation processes.

## Introduction

Crassulacean acid metabolism (CAM) species exhibit rescheduled diel opening of stomata, enabling uptake of atmospheric CO_2_ during the cooler, more humid conditions at night. Stomatal closure during the hotter, less humid day-time hours minimizes water loss through transpiration, thereby facilitating survival in water-limited environments (Borland et al., 2009). The diel cycle of CAM photosynthesis is generally delineated into four distinct phases representing characteristic patterns of gas exchange and fluctuations in the abundance of key metabolites that define carbon supply and demand over the 24-h cycle (Osmond et al., 1978). At night, atmospheric CO_2_ is fixed to phospho*enol*pyruvate (PEP), which is provided via glycolytic breakdown of storage carbohydrates (i.e. soluble sugars or starch) accumulated during the previous day, by the enzyme phospho*enol*pyruvate carboxylase (PEPC) leading to nocturnal accumulation of malic acid in the vacuole (Phase I). During the light period, PEPC is deactivated and malic acid is released from the vacuole for its decarboxylation either by malic enzyme (ME) or phospho*enol*pyruvate carboxykinase (PEPCK), depending on the species, behind shut stomata (Dittrich, 1976; Holtum et al., 2005). Liberated CO_2_ accumulates within the leaf tissue and is refixed by ribulose-1,5-bisphosphate carboxylase/oxygenase (Rubisco) in the Calvin-Benson-Bassham (CBB) cycle (Phase III). These major CAM phases are flanked by Phase II at the start of the day and Phase IV at the end of the day, when stomata gradually close and re-open respectively (Borland and Taybi, 2004). The four CAM phases exhibit significant plasticity in duration and magnitude, which is a key factor to improve carbon gain and water use under changing environmental conditions (Lin and Hsu, 2004; Ceusters et al., 2011; Tay et al., 2019; Ceusters et al., 2021b).

In ME-type CAM plants, diurnal liberation of CO_2_ from malate can be catalyzed either by mitochondrial NAD-ME and/or cytosolic/plastidic NADP-ME, yielding pyruvate. RNA interference (RNAi) silencing of *NAD-ME* (β-subunit) in *Kalanchoë fedtschenkoi* determined NAD-ME as the main decarboxylase in this species, with only a minor contribution from NADP-ME (Dever et al., 2015). Reducing NAD-ME activity also led to the loss of rhythmic oscillations of core circadian clock transcripts (*KfTOC1*), indicating that disrupting the decarboxylation process of CAM may feedback to affect the central circadian clock (Dever et al., 2015). The *α* and *β* subunits of plant mitochondrial NAD-ME are each encoded by an individual gene and their expression is tied to the central circadian clock (Dever et al., 2015; Yang et al., 2017; Francisco et al., 2021). A level of transcriptional control over the day-time breakdown of malate via ME is also suggested in *K. fedtschenkoi*, with maximal *Kf*NAD ME1 protein abundance noted in the middle of the photoperiod at the time when malate decarboxylation rates are maximal (Abraham et al., 2020). Moreover, ME-type CAM species require cytosolic/plastidic pyruvate orthophosphate dikinase (PPDK) to convert ME-derived pyruvate to PEP, initiating the gluconeogenic biosynthesis of starch. Its activity in the cytosol is reported to be twice as high compared to its chloroplastic counterpart in *K. fedtschenkoi* (Kondo et al., 2000). In C_3_ and C_4_ plants, light-dependent inactivation-activation of PPDK is mediated by a bifunctional kinase/phosphatase PPDK-regulatory protein (RP) that catalyzes reversible phosphorylation (inactive form) and dephosphorylation (active form) of PPDK in the dark and light, respectively (Slack, 1968; Burnell and Hatch, 1985; Chastain et al., 2002; Chastain and Chollet, 2003; Astley et al., 2011). Dever et al. (2015) reported that this light/dark phospho-regulation of PPDK activity during the diel cycle also occurs in *K. fedtschenkoi*, similar to the chloroplastic forms in C_3_ and C_4_ plants. Silencing of *PPDK* also affected major CAM components such as NAD-ME and PEPC, which was reflected at the transcript, protein, and enzyme activity levels (Dever et al., 2015). Abraham et al. (2020) reported diel changes in phosphopeptide abundance of PPDK protein in *K. fedtschenkoi* and indicated phosphorylation during the night. Additionally, an orthologue of PPDK-RP showed increased protein abundance at dusk and for most of the night (Abraham et al., 2020).

Whilst the nocturnal processes of primary carbon fixation, malate biosynthesis and its transport into the vacuole are well known, less information exists about the diurnal biochemical process of malate remobilisation and its regulation during the CAM cycle (Ceusters et al., 2021a; Winter and Smith, 2022). It has long been questionable whether malate efflux from the vacuole, the enzymes responsible for malate decarboxylation, or the direct light-dependent assimilation of CO_2_ via Rubisco in the chloroplasts set the pace for diurnal processing of accumulated malate (Lüttge, 2002). Supported by detailed stoichiometric analyses based on existing literature, Ceusters et al. (2021) postulated that the vacuolar efflux of malate itself might be the primary candidate determining the rate/duration and the onset of diurnal deacidification in CAM leaves. Intrinsic activities of NAD(P)-ME, PEPCK (CO_2_ liberation) and Rubisco (CO_2_ uptake) in leaf mesophyll cells during Phase III, across various CAM species under normal homeostatic conditions, have been found to be far in excess of those needed to accommodate the observed rates of malate degradation.

Malate efflux from the vacuole remains one of the least understood processes in CAM research, with proposed mechanisms including passive diffusion, a proton-linked symporter and/or a dedicated malate anion channel (Smith et al., 1996). Passive diffusion of malic acid out of the vacuole is relevant only at the lowest vacuolar pH around dawn (Lüttge and Smith, 1984). For most of the light period, vacuolar efflux mainly occurs as Hmalate^-^ and or malate^2-^, with substantial evidence indicating an intimate stoichiometry of 2H^+^:malate^2-^ (or alternatively 1H^+^:Hmalate^-^). Electrophysiological measurements in *Kalanchoë daigremontiana* unveiled a strongly inward-rectifying malic acid-selective anion channel in CAM plant tonoplasts responsible for malate influx into the vacuole (Hafke et al., 2003). Whilst the diurnal malate tonoplast exporter remains unidentified, ALUMINIUM-ACTIVATED MALATE TRANSPORTER 9 (ALMT9) has emerged as the molecular candidate for the tonoplast malate influx channel at night, given its observed increase in transcript levels during the transition from C_3_ to CAM (Brilhaus et al., 2016). This strongly rectifying channel, importing malate, could also serve as a potential export pathway if a reverse electrochemical gradient is assumed or if posttranslational modifications alter its transport properties. Tonoplast-localized members of the *ALMT* gene family and the tonoplast dicarboxylate transporter (tDT) have been put to front as potential candidates for facilitating the diurnal release of malate from the vacuole in CAM plants and have recently been discussed by Ceusters et al. (2021). While tDT is encoded by a single gene in all plant genomes sequenced to date (Wai et al., 2017), the *ALMT* gene family in Arabidopsis consists of 14 members (Hoekenga et al., 2006). Each of these members exhibits specific subcellular localization (either the plasma membrane or the tonoplast), cell type specificity (root, shoot, mesophyll and/or guard cells), and transport direction (facilitating malate efflux, influx, or both), with approximately half of the members being functionally well characterized in Arabidopsis (Doireau et al., 2024). Electrophysiological approaches in Arabidopsis showed that *At*tDT facilitates vacuolar malate fluxes in both directions in exchange for cytosolic citrate (Frei et al., 2018) and that *At*ALMT4 can mediate vacuolar malate efflux in guard cells during ABA-induced stomatal closure (Eisenach et al., 2017). Temporal diel transcript abundance data of various (facultative) CAM species suggest a specific role for both putative malate exporters in CAM functioning (Brilhaus et al., 2016; Wai et al., 2017; Yang et al., 2017; Ferrari et al., 2020; Zhang et al., 2020; Gilman et al., 2022).

With regard to environmental factors affecting diurnal malate remobilisation, an important role for the photosynthetic photon flux density (PPFD) has already been acknowledged for a long time. Different observations have indicated that PPFD determines the rate of organic acid mobilization from the vacuole (Kluge, 1968; Barrow and Cockburn, 1982; Thomas et al., 1987), which seems to be connected to the rates of electron transport and Rubisco-mediated CO_2_ uptake *in vivo* (Lüttge, 2004). In leaves of *K. fedtschenkoi* exposed to continuous darkness, high background malic acid levels have been reported (Anderson and Wilkins, 1989). These findings suggest that specific environmental conditions can lead to uncoupling between malate efflux and its further processing, supporting the view that the efflux process is likely governed by an integration of circadian and metabolic signals (Ceusters et al., 2021a). Research on the influence of photoperiod on CAM has primarily focused on the free running circadian rhythms under continuous light and dark conditions (Hartwell et al., 1996; Dodd et al., 2003; Davies and Griffiths, 2012) and on photoperiodic control of CAM expression in *Kalanchoë blossfeldiana* and *Mesembryanthemum crystallinum* under short and long days (Brulfert et al, 1975; Brulfert et al., 1982; Cheng and Edwards, 1991). However, the precise impact of light intensity and photoperiod on the molecular mechanisms underlying malate transport and processing – such as putative tonoplast malate exporters and the enzymes involved in malate remobilisation – have received marginal attention in CAM research.

The present work aims to provide mechanistic insights into the regulation of light intensity and photoperiod (single-day treatment) on different aspects related to the diurnal process of malate remobilisation from the vacuole in CAM mesophyll cells. Employing an integrative analysis that spans physiological, temporal metabolic, protein, enzymatic, and gene transcript levels, this study particularly focusses on (1) the release of malate from the vacuole via either ALMT and/or tDT, (2) malate decarboxylation by ME, (3) refixation of CO_2_ by Rubisco and recycling of ME-derived pyruvate to starch via PPDK.

## Materials and Methods

### Plant material and growth conditions

*Kalanchoë fedtschenkoi* Hamet et Perrier plants were propagated clonally from leaf margin adventitious plantlets and were grown grouped by four in 3.4 L pots containing universal potting soil (De Ceuster Meststoffen NV, Belgium). Plants were cultivated for 10-12 weeks in controlled growth rooms with a photoperiod of 12 h (from 08:00 until 20:00 h), day/night temperature of 23/18 °C, and photosynthetically active radiation (PAR) at the apex of ∼300 µmol m^-2^ s^-1^ which was provided by LED growth lamps with broad wavelength spectra (Mars Hydro TSL-2000: https://marshydro.eu/) until reaching the 12-leaf pairs (LP) stage. Watering was performed by pouring twice a week and plants were maintained under ambient CO_2_ concentrations.

### Experimental design and temporal sampling

A first group of plants of the obligate CAM model species *K. fedtschenkoi*, with 12 leaf pairs, was exposed to a range of decreasing light intensities [i.e. 300 (control)-200-100-50-10 µmol m^-2^ s^-1^] during a single light period to determine the impact of light-limiting conditions on the rate/duration of diurnal malate processing (light intensity experiment). Light intensities were checked using a handheld full-spectrum quantum meter (Apogee MQ-501). A second group of plants was subjected to different photoperiodic treatments to examine the effect of photoperiod on malate processing (photoperiod experiment). Lights were switched on at 08:00 h for the control treatment (12L/12D), at 04:00 h for the extended photoperiod (16L/8D), at 12:00 h for the curtailed photoperiod (8L/16D), and did not switch on for the continuous dark treatment (24D). Separate but uniform batches of plants were used for both experiments. Samples were taken from leaf pairs 5, 6, and 7 (LP5, 6, 7; with LP1 starting from apex) in a 2-h time interval during the light period, starting 15 min before the onset of the light period (i.e. 7:45 h and 3:45 h for the light intensity and photoperiod experiment respectively) and ending 15 min before the end of the light period (i.e. 19:45 h for both experiments). Five biological replicates were taken at each timepoint and were snap frozen in liquid nitrogen, powdered and stored at −80 °C until further analysis.

### Leaf CO_2_ exchange measurements and biochemical analyses of metabolites

Leaf CO_2_ exchange data was collected randomly on leaf pairs 5, 6, and 7. Leaves were placed into a broad leaf chamber (6.25 cm^2^) of the LCi Portable Photosynthesis System (ADC BioScientific Ltd., United Kingdom) 1 h before the start of the photoperiod. The incoming air was passed through a 20-l bottle to buffer short-term fluctuations in the CO_2_ concentration. Measurements were recorded every 15 min. Each gas exchange curve presented is representative of data obtained from three independent biological replicates.

Metabolite analyses were performed on a 2-h time resolution (n=5). Determination of starch content was performed as described in Methods S1. Extraction and measurement of malic acid was performed as described by Chen et al. (2002), but with modifications as described in Methods S1. In the photoperiod experiment, sucrose content was determined using an enzyme-coupled spectrophotometric assay according to the manufacturer’s protocol (Megazyme, K-SUFRG), to correlate with starch concentrations (Methods S1).

### Enzyme activity assays

Enzyme activity analyses were performed on a 4-h time resolution. Extraction and assay for mitochondrial NAD-ME, cytosolic/plastidic NADP-ME, and PPDK were based on the methods described by Dever et al. (2015), with slight modifications as described in Methods S1. The extraction and assay of Rubisco were based on the method described by Borland et al. (1998), with slight modifications as described in Methods S1.

### Western blotting

PPDK protein abundance was determined via Western blotting using specific antibody from Agrisera (Vännäs, Sweden) (AS13 2647), following general practices as described in more detail in Methods S1. Phosphorylated PPDK (P-PPDK) antibody [raised against maize (*Zea mays*) PPDK] was kindly provided by Chris J. Chastain, Minnesota State University, Moorhead (Chastain et al., 2018).

### RNA extraction, RNA sequencing, and bioinformatics

A selection of light treatments and timepoints was made prior to performing the time-course bulk RNA-seq experiment. For the light intensity experiment, treatments of 300-200-100 µmol m^-2^ s^-1^ and timepoints 08:00 h, 14:00 h, and 20:00 h were selected. In the photoperiod experiment, all treatments were analyzed, and samples were taken from timepoints 06:00 h, 10:00 h, 14:00 h, and 18:00 h (i.e. 2 h after lights were switched on in a treatment). Total RNA was extracted from three biological replicates using the Qiagen RNeasy Plant Mini Kit following the manufacturer’s protocol, with modifications described in Methods S1. Genomic DNA was removed via on-column DNase digestion (Qiagen). RNA quantity, quality, and integrity were assessed by nanodrop (NanoDrop Lite Plus Spectrophotometer, Thermo Scientific), gel electrophoresis, and on the Bioanalyzer (Agilent) respectively. Library preparation and sequencing were performed at Azenta (Genewiz) using the Illumina NovaSeq platform (2×150 bp paired-end sequencing) to obtain 17-55 million reads per sample. Considering the primary narrative of the manuscript, the RNA-seq data analysis was exclusively centered on genes associated with diurnal malate remobilisation. Further information regarding RNA-seq data analysis is provided in Methods S1.

### Data analysis

Data analysis was performed using SPSS 28.0 (IBM, New York, NY, USA). Before carrying out statistical tests, normality and equality of variances of the data were checked by means of a Shapiro-Wilk and Levene’s test (p>0.05), respectively. Means of two groups were compared by a two sample t-test. Means of three or more groups were compared by ANOVA followed by Tukey’s HSD post-hoc test (p<0.05). The non-parametric Kruskal-Wallis test, followed by Dunn’s post-hoc test, was used if the conditions were not met.

## Results

### Effects of light intensity and photoperiod on the magnitude and the duration of the different CAM phases of diel CO_2_ exchange

Control plants exposed to 300 µmol m^-2^ s^-1^ exhibited a typical CAM pattern with nocturnal CO_2_ fixation (Phase I), net CO_2_ loss during the day (Phase III), and two intermediate phases (Phases II and IV) (Fig. 1a). A reduction in light intensity to 200 µmol m^-2^ s^-1^ caused a prolongment of Phase III and resulted in a substantial drop in both the amplitude and duration of Rubisco-mediated net CO_2_ uptake during Phase IV. Nocturnal carboxylation was less affected as these plants still fixed about 75% compared to controls. Under lower light intensities (100, 50 or 10 µmol m^-2^ s^-1^) Phase IV was completely abolished and below 100 µmol m^-2^ s^-1^ during the light period, no net nocturnal carboxylation was observed anymore (Fig. 1a).

**Figure 1.**
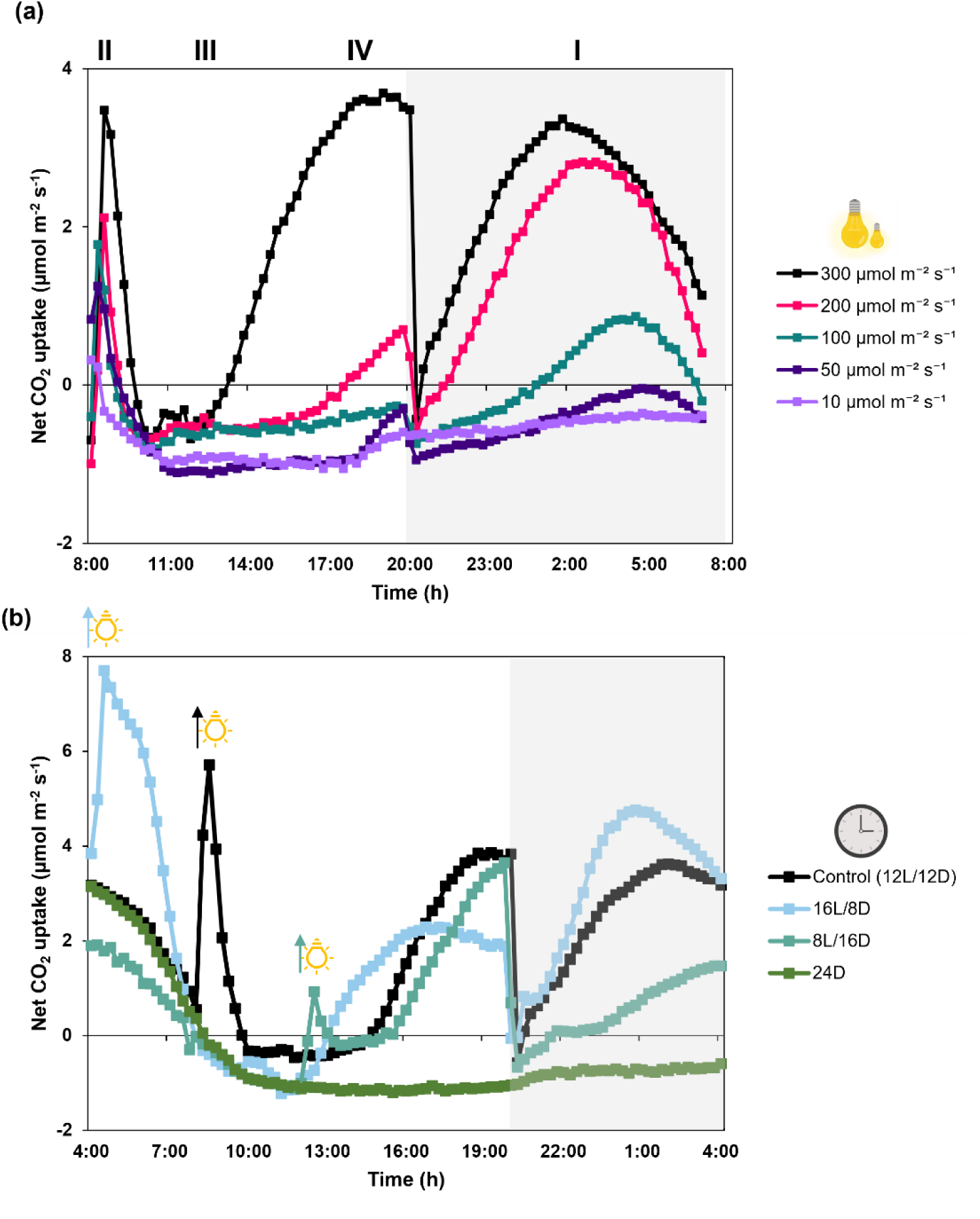
Diel leaf CO_2_ uptake measurements for *Kalanchoë fedtschenkoi* under different light intensity and photoperiodic conditions. Net 24-h CO_2_ uptake (µmol m^-2^ s^-1^) measured for leaf pairs 5, 6, or 7 of *K. fedtschenkoi* exposed to reduced light intensity (a) and different photoperiodic (b) conditions for one day. Roman numbers above Fig. 1a indicate the different CAM phases. The dark period is indicated in grey in both figure panels, while arrows with a lamp icon indicate the onset of light for the corresponding photoperiod treatment in Fig. 1b. Gas exchange curves are representative of three replicate runs with SE < 15% (n=3 plants).

Plants used in the photoperiod experiment also showed a typical CAM pattern under control photoperiodic conditions (i.e. 12L/12D) (Fig. 1b). Extending the photoperiod by switching lights on at 04:00 h (16L/8D) caused a substantial rise in net carbon uptake between 04:00 h and 08:00 h (Phase II), attributed to the combined action of PEPC and Rubisco (Fig. 1b). The extended photoperiod also led to a curtailment of Phase III and an improved nocturnal carboxylation (+33%) between 20:00 and 04:00 h compared to controls. Plants exposed to a shortened photoperiod, with lights switched on at 12:00 h (8L/16D), showed only a short negligible Phase II and a curtailed Phase III whilst Phase IV was remarkably similar in magnitude and duration to those of controls. Nocturnal CO_2_ sequestration was also clearly diminished by the shortened photoperiod. Under continuous darkness (24D), plants exhibited continuous respiratory release of CO_2_ throughout the diel cycle, starting at 08:00 h.

### Effects of light intensity and photoperiod on the core biochemical aspects of diel CO_2_ fixation

As expected, large differences in malate levels were measured between dawn (08:00 h) and dusk (20:00 h) (ca. 30 µmol malate g^-1^FW) in leaves exposed to high light intensities of 300 and 200 µmol m^-2^ s^-1^ (Fig. 2a). Typically, starch showed an opposite trend compared to malate, accumulating to high levels (ca. 40 µmol Glc eq. g^-1^FW) at dusk (20:00 h) in these leaves (Fig. 2b). Leaves exposed to 100 µmol m^-2^ s^-1^ showed a moderate acid degradation (ca. 20 µmol malate g^-1^FW), whereas little to no malate degradation was observed at 50 and 10 µmol m^-2^ s^-1^. Accordingly, starch content at dusk was strongly reduced at 100, 50, and 10 µmol m^-2^ s^-1^ (ca. 14 µmol Glc eq. g^-1^FW).

**Figure 2.**
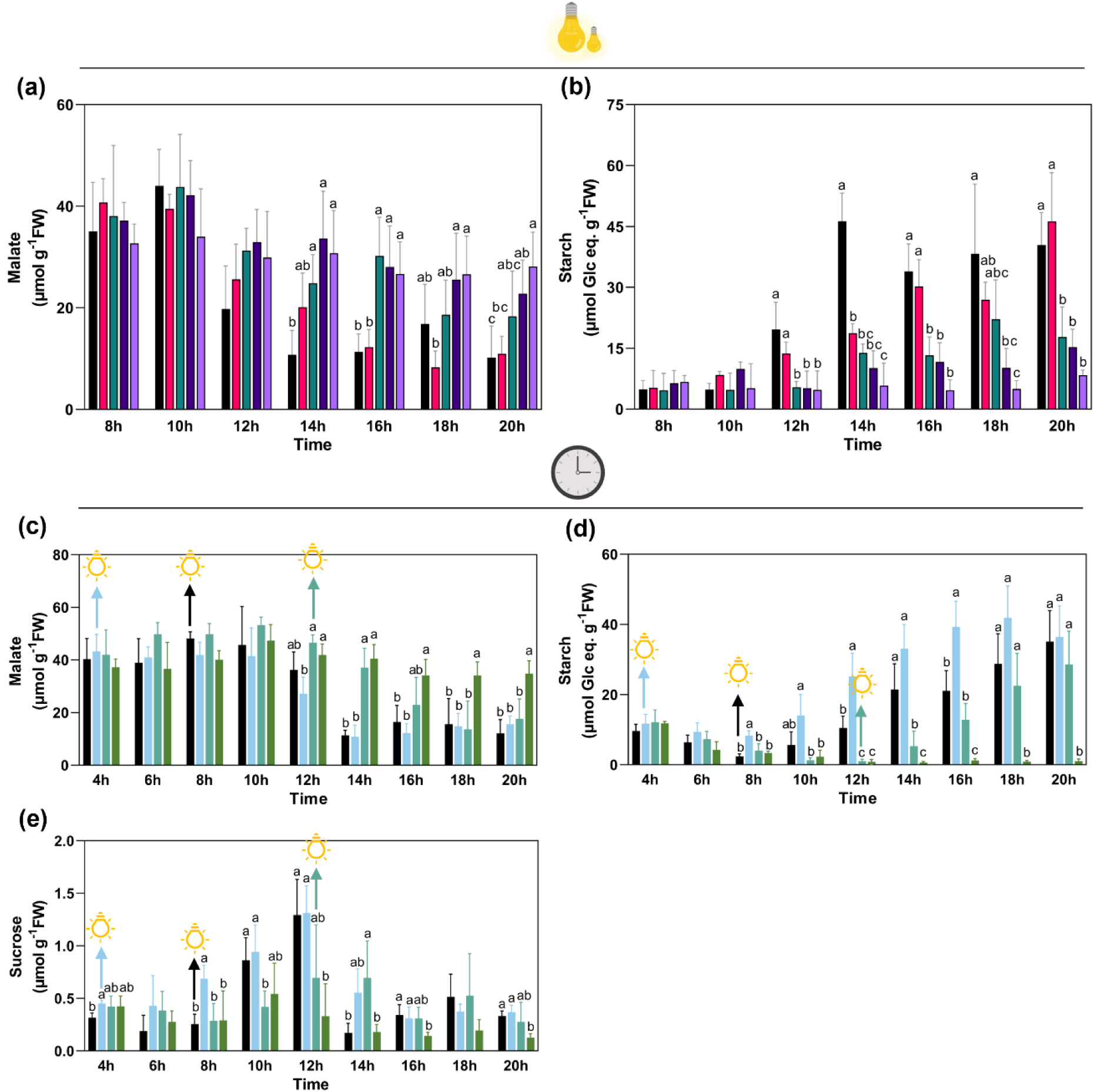
Effects of light intensity and photoperiod on the temporal pattern of key CAM metabolite concentrations. Time course metabolite data for malate (µmol g^-1^FW) (a) and starch (µmol Glc eq. g^-1^FW) (b) between 08:00h (dawn) and 20:00h (dusk) in the light intensity experiment. Time course metabolite data for malate (µmol g^-1^FW) (c), starch (µmol Glc eq. g^-1^FW) (d), and sucrose (µmol g^-1^FW) (e) between 04:00h and 20:00h in the photoperiod experiment. Metabolites were measured randomly for leaf pairs 5, 6, or 7 of *Kalanchoë fedtschenkoi* exposed to reduced light intensities and different photoperiodic conditions for one day. Arrows with a lamp icon indicate the onset of light for the corresponding photoperiod treatment. Data are means ± SE (n=5 plants). Values were compared among the different treatments per time point according to Tukey’s HSD test at p<0.05 marked by different letters.

Slopes of linear curves fitted to the data points (malate and starch) at timepoints 10:00, 12:00, 14:00, and 16:00 h (CAM Phase III) were determined and served as a proxy for malate degradation and starch accumulation rates (Table S1). Plants at 300 µmol m^-2^ s^-1^ exhibited the highest rate of acid degradation and starch accumulation, which led to the malate and starch pool reaching their minimum and maximum levels sooner (around 14:00 h) compared to treatments with reduced light intensity (Figs 2a, b). Generally, the rates of malate degradation and starch accumulation gradually declined with decreasing light intensities, with significant lower rates (p<0.05) particularly noticed from 100 µmol m^-2^ s^-1^ onwards (for starch significantly lower rates were already noticed at 200 µmol m^-2^ s^-1^ (p<0.05)) (Table S1).

Remarkably, in both the extended (16L/8D) and control (12L/12D) photoperiodic treatments, degradation of malate was initiated around 10:00 h (p<0.05). This observation indicates that an earlier start of the photoperiod at 04:00 h did not induce a premature malate degradation (Fig. 2c). However, the onset of the lights at 04:00 h slowed down the remaining starch degradation (p<0.05 at 08:00 h), increased sucrose content at 08:00 h (p<0.05), and brought about an increased rate of starch accumulation during the photoperiod (Figs 2d, e). Under an extended dark period (8L/16D), starch was completely exhausted by 10:00 h and malate levels remained high until 2 h after the onset of the lights (at 14:00 h). However, the remaining photoperiod seemed still sufficient to allow both malate and starch levels to reach equal values compared to the extended and control photoperiodic treatments before dusk (18:00 h, p>0.05) (Figs 2c, d). Under continuous darkness (24D), malate levels remained high comparable to dawn levels in controls whilst starch concentrations reached near-zero levels from 10 h.

### Intrinsic enzyme activities of NAD-ME, Rubisco and PPDK but not NADP-ME respond to light intensity and photoperiod

At dawn, relatively high activity levels were measured for mitochondrial NAD-ME (12 µmol PYR h^-1^ g^-1^FW), whilst cytosolic/plastidial NADP-ME activity was much lower (approximately 2.5 µmol PYR h^-1^ g^-1^FW) (Figs 3a, b, e, f). At high light intensities (300 and 200 µmol m^-2^ s^-1^), NAD-ME activity almost doubled during the latter half of the light period to ca. 20 µmol PYR h^-1^ g^-1^FW (Fig. 3a). At lower light intensities (100, 50, and 10 µmol m^-2^ s^-1^), NAD-ME retained its baseline activity throughout the light period (Fig. 3a). The observed increase in NAD-ME activity – doubling its activity levels and peaking at around 25 µmol PYR h^-1^ g^-1^FW - was ahead under an extended (16L/8D) photoperiod and lagged behind under a curtailed (8L/16D) photoperiod compared to controls. Under continuous dark conditions, NAD-ME activity remained at baseline levels during the complete sampling period (Fig. 3e). In contrast to the time- and light-responsive nature of NAD-ME, NADP-ME activity levels remained fairly constant throughout the light period across all timepoints and light treatments (Figs 3b, f).

**Figure 3.**
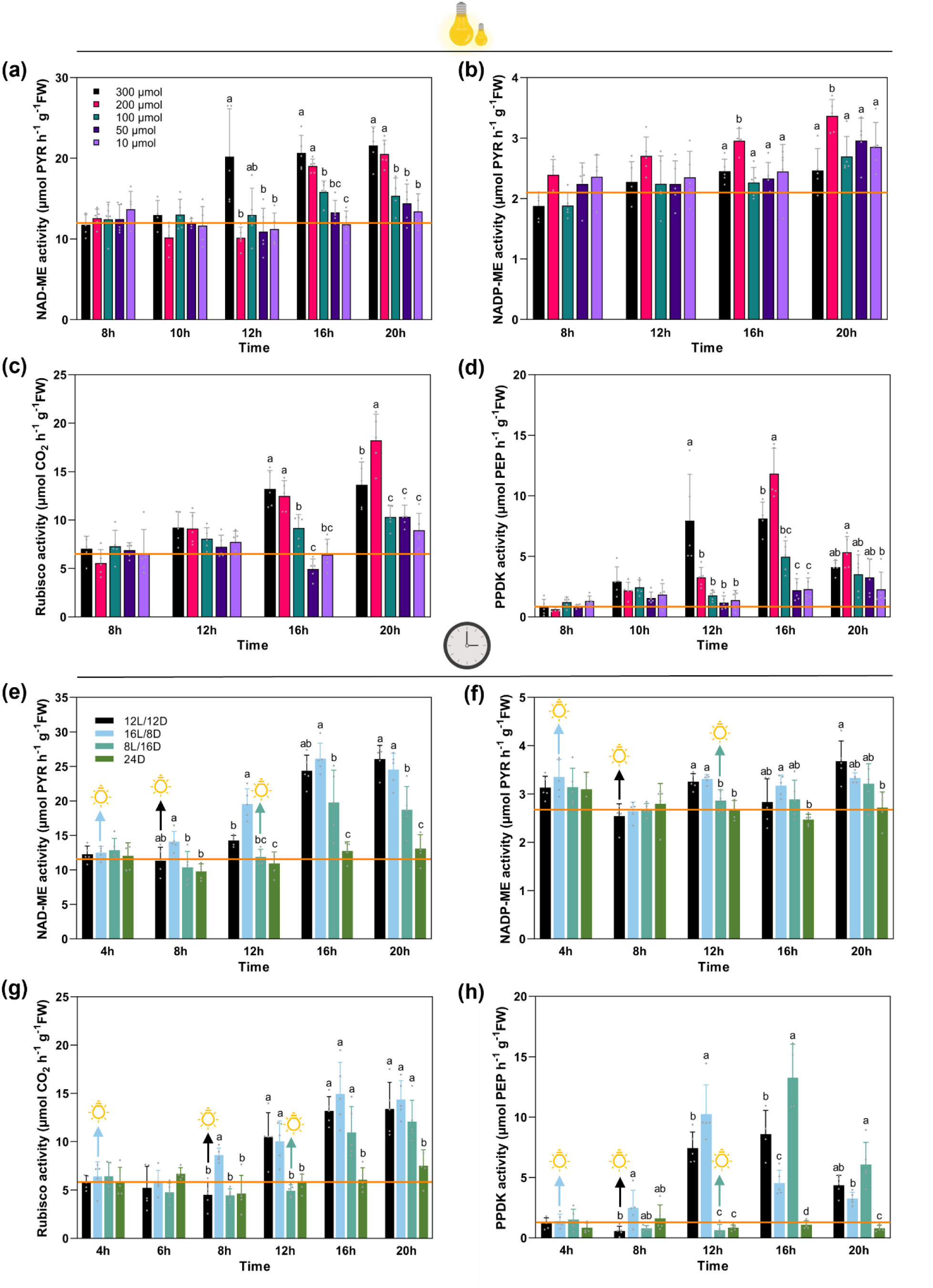
Temporal intrinsic activity data of enzymes potentially involved in diurnal malate remobilisation from the vacuole in CAM. Time course enzyme activity data for mitochondrial NAD-ME (µmol PYR h^-1^ g^-1^FW) (a), cytosolic/chloroplastic NADP-ME (µmol PYR h^-1^ g^-1^FW) (b), Rubisco (µmol CO_2_ h^-1^ g^-1^FW) (c), and PPDK (µmol PEP h^-1^ g^-1^FW) (d) between 08:00h and 20:00h measured for leaf pairs 5, 6, or 7 of *Kalanchoë fedtschenkoi* exposed to different light intensities for one day. Time course enzyme activity data for NAD-ME (e), NADP-ME (f), Rubisco (g), and PPDK (h) between 04:00h and 20:00h measured for leaf pairs 5, 6, or 7 of *K. fedtschenkoi* exposed to different photoperiodic conditions for one day. NAD-ME and PPDK activity was also measured at 10:00 h in the light intensity experiment to determine whether the observed increase already occurred before 12:00 h. Rubisco activity was also measured at 06:00 h in the photoperiod experiment to determine whether the observed increase in Rubisco-mediated carbon uptake between 04:00 and 08:00 h under an extended photoperiod (16L/8D) (Fig. 1b) was also reflected in intrinsic Rubisco activity. Orange lines indicate base levels of enzyme activity at dawn. Arrows with a lamp icon indicate the onset of light for the corresponding photoperiod treatment. Data are means ± SE (n=5 plants). Values were compared among the different treatments per time point according to Tukey’s HSD test at p<0.05 marked by different letters.

Rubisco activity displayed a baseline level of about 6.5 µmol CO_2_ h^-1^ g^-1^FW at dawn (Figs 3c, g). Under high light intensities (300 and 200 µmol m^-2^ s^-1^), Rubisco activity nearly doubled, reaching levels about 13 µmol CO_2_ h^-1^ g^-1^FW in the latter half of the light period (Fig. 3c). This increase occurred earlier under an extended (16L/8D) photoperiod and later under a curtailed (8L/16D) photoperiod compared to controls. In continuous darkness, Rubisco activity was maintained at baseline levels (Fig. 3g).

PPDK exhibited a relatively low baseline activity of approximately 1 µmol PEP h^-1^ g^-1^FW at dawn, reaching a peak around midday/afternoon (ca. 9 µmol PEP h^-1^ g^-1^FW) under high light intensities, before declining again towards dusk (Fig. 3d). The increase in PPDK activity, up to ninefold, occurred sooner under an extended (16L/8D) photoperiod and later under a curtailed (8L/16D) photoperiod compared to controls. Consequently, lower activity levels at dusk were also reached sooner and later, respectively (Fig. 3h). Notably, under the shortened photoperiod, PPDK activity peaked at 13.2 µmol PEP h^-1^ g^-1^FW at 16:00 h, exceeding the maximum levels observed at 12:00 h in leaves under the control and extended photoperiod treatments. A significant increase/decrease pattern was absent in leaves exposed to low light intensities (50 and 10 µmol m^-2^ s^-1^) and continuous darkness, where PPDK activity remained close to low baseline levels throughout the light period (Figs 3d, h).

Given the strong light sensitivity of PPDK activity, protein abundances for both PPDK (active form) and P-PPDK (inactive form) were also determined. Under control conditions (300 µmol m^-2^ s^-1^ and 12L/12D) protein abundances of both PPDK and P-PPDK were increasing throughout the photoperiod but especially the latter showed a higher increase (Figs 4a, b). A reduced light intensity of 100 µmol m^-2^ s^-1^ resulted in a declined protein abundance of PPDK, whereas P-PPDK abundance was slightly higher at 12:00 h and similar for the timepoints of 16:00 and 20:00 h compared to controls (Fig. 4a). This pattern also aligns with measured PPDK activity at 12:00 h, which was highest at 300 µmol m^-2^ s^-1^ (high PPDK and low P-PPDK abundance) and significantly lower at 100 µmol m^-2^ s^-1^ (low PPDK and high P-PPDK abundance). At 04:00 h, prior to the onset of photoperiodic treatments, both PPDK and P-PPDK protein levels were relatively high compared to dawn (08:00 h) for all treatments (Fig. 4b). For the extended photoperiod (16L/8D), the abundance of both PPDK and P-PPDK was already higher at 12:00 h compared to the other photoperiodic treatments. At 16:00 h in the photoperiod experiment, PPDK activity peaked under the shortened photoperiod (8L/16D), while Westerns also showed the highest PPDK abundance and lowest P-PPDK abundance compared to other treatments at this timepoint (Fig. 4b), further illustrating the relationship between PPDK activity and the abundance of active PPDK and inactive P-PPDK. Under continuous darkness, the levels of both PPDK and P-PPDK protein remained remarkably lower at each timepoint (Fig. 4b).

**Figure 4.**
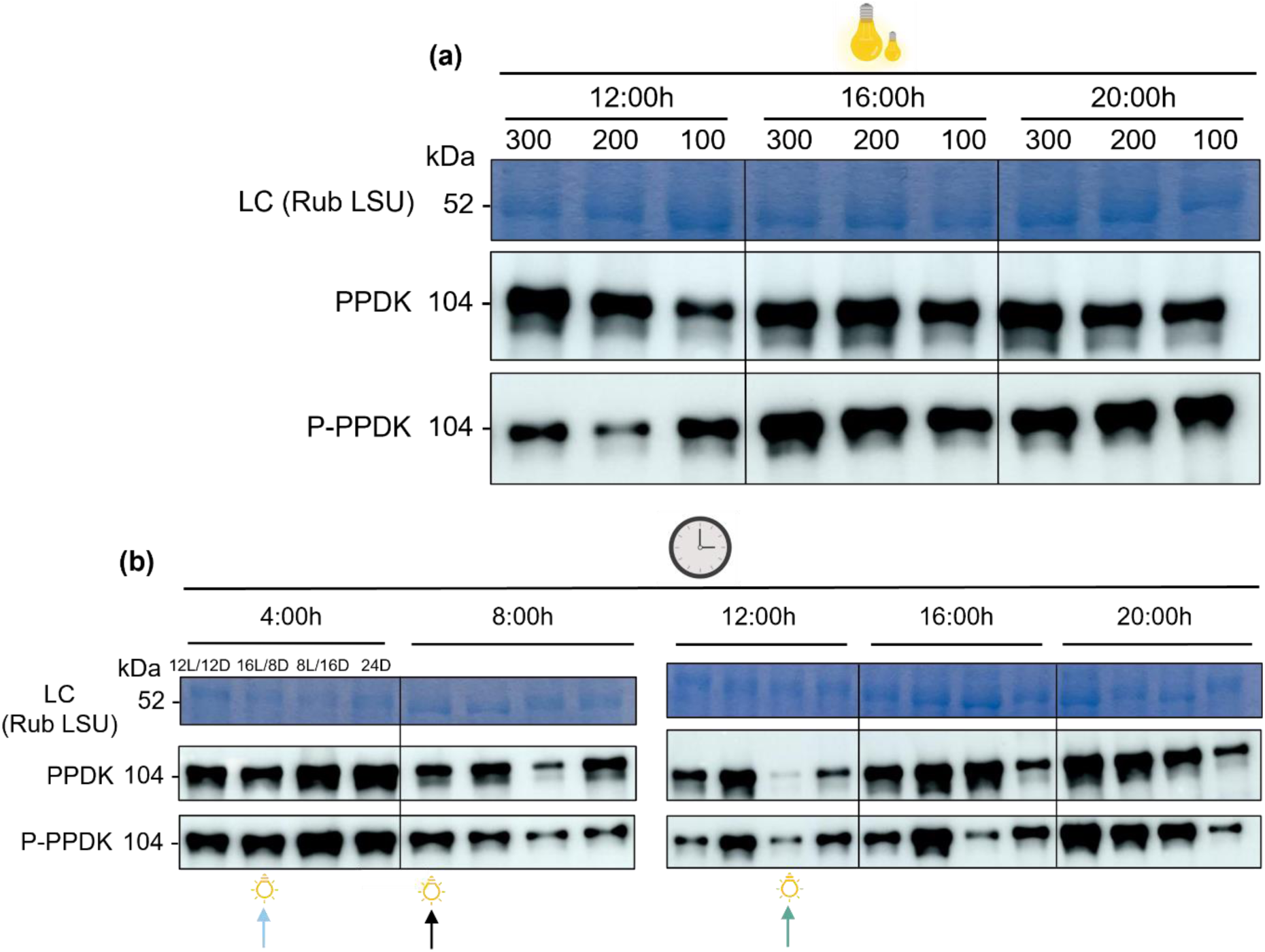
Temporal protein abundance pattern of PPDK and P-PPDK under different light intensities and photoperiods. Time course Western blots showing the abundance of PPDK and P-PPDK protein at 12:00h, 16:00h and 20:00h measured for leaf pairs 5, 6, or 7 of *Kalanchoë fedtschenkoi* exposed to different light intensities for one day (a). Time course Western blots showing the abundance of PPDK and P-PPDK protein between 04:00h and 20:00h measured for leaf pairs 5, 6, or 7 of *K. fedtschenkoi* exposed to different photoperiodic conditions for one day (b). The Rubisco large subunit (LSU) band on a stained Coomassie gel served as a loading control (LC). Arrows with a lamp icon indicate the onset of light for the corresponding photoperiod treatment.

### RNA sequencing uncovers light-responsive and temporal transcript abundance patterns of genes involved in diurnal malate remobilisation

Our time-course RNA-seq analysis provided insights into potential light-responsive transcript abundance patterns of genes encoding proteins putatively involved in diurnal malate remobilisation in CAM, including ALMT, tDT, NAD(P)-ME, Rubisco, Rubisco activase, PPDK, and PPDK-RP. Heatmaps were created to highlight which genes in *K. fedtschenkoi* leaves were responsive to changes in light intensity and photoperiod (Fig. 5). For gene families containing multiple members, only the two to three genes with the highest transcript abundance (considered to be the functional members of their gene families) were selected and depicted in the heatmaps (Fig. 5). At the start of each section below, the total number of genes in the specific gene family is specified. Since these heatmaps do not convey the temporal patterns of transcript abundance during the light period across all tested light conditions, bar charts depicting the average normalized counts for all light treatments, sampling timepoints, and gene family members were provided in Supporting Information Figs S1-S6.

**Figure 5.**
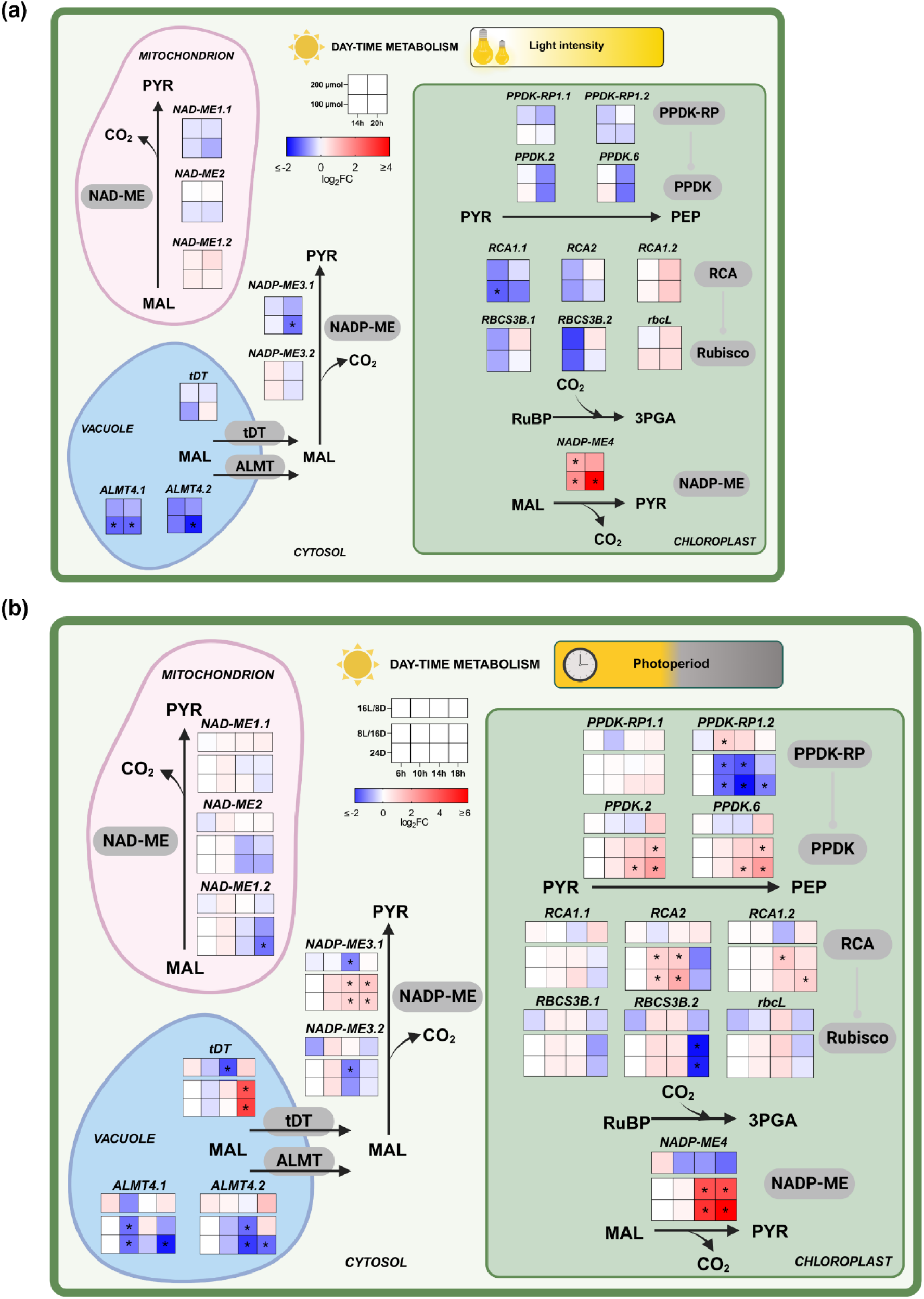
Light-induced transcript level changes of key genes potentially involved in diurnal malate remobilisation in leaves of *Kalanchoë fedtschenkoi*. Heatmaps indicate log_2_(fold-change) of transcript abundance (blue represents downregulation, red represents upregulation) in leaves under different light intensities (a) and photoperiods (b) at different diurnal time points. The legend is depicted in the upper middle of the figures. Statistical differential gene expression analysis was done with DESeq2. Significantly differential expression in comparison with 300 µmol m^-2^ s^-1^ (a) or 12L/12D photoperiod (b) is indicated by an asterisk (adjusted p-value < 0.05, |log_2_FC|>1) in each box. Three biological replicates were used for RNA-seq analysis. Black arrows represent core reactions in diurnal malate remobilisation whereas gray lines terminated by a closed circle indicate regulatory interactions. Enzymes are presented by gray ovals. Bar charts depicting average normalized counts for each treatment at each time point, and all gene family members, are shown in Supporting Information Figs S1-S6. Metabolites: 3PGA, 3-phosphoglycerate; MAL, malate; PEP, phospho*enol*pyruvate; PYR, pyruvate; RuBP, ribulose 1,5-bisphosphate. Enzymes: ALMT, tDT, NAD-ME, NADP-ME, PPDK, PPDK-RP, RCA, Rubisco.

#### Vacuole

The *ALMT* gene family contains 13 members in *K. fedtschenkoi*. Two gene orthologs to *AtALMT4* (At1g25480) [*ALMT4.1* (Kaladp0024s0194) and *ALMT4.2* (Kaladp0073s0021), with the latter having the highest transcript abundance of both] were the most highly expressed members of the *ALMT* gene family in *K. fedtschenkoi* leaves and were thus considered as putative functional candidates to potentially accommodate vacuolar malate efflux (Figs 6a, b, S1). Under control conditions (300 µmol m^-2^ s^-1^ and 12L/12D), both *ALMT4* genes displayed a temporal transcript abundance pattern with relatively low transcript levels at dawn and higher transcript abundance during the diurnal period (Figs 6a, b). Under light-limited conditions (lower light intensities, shortened photoperiod and continuous darkness), transcript abundances of both *ALMT4* genes were generally lower throughout the light period than those observed in control light treatments (Fig. 5, Figs 6a, b).

**Figure 6.**
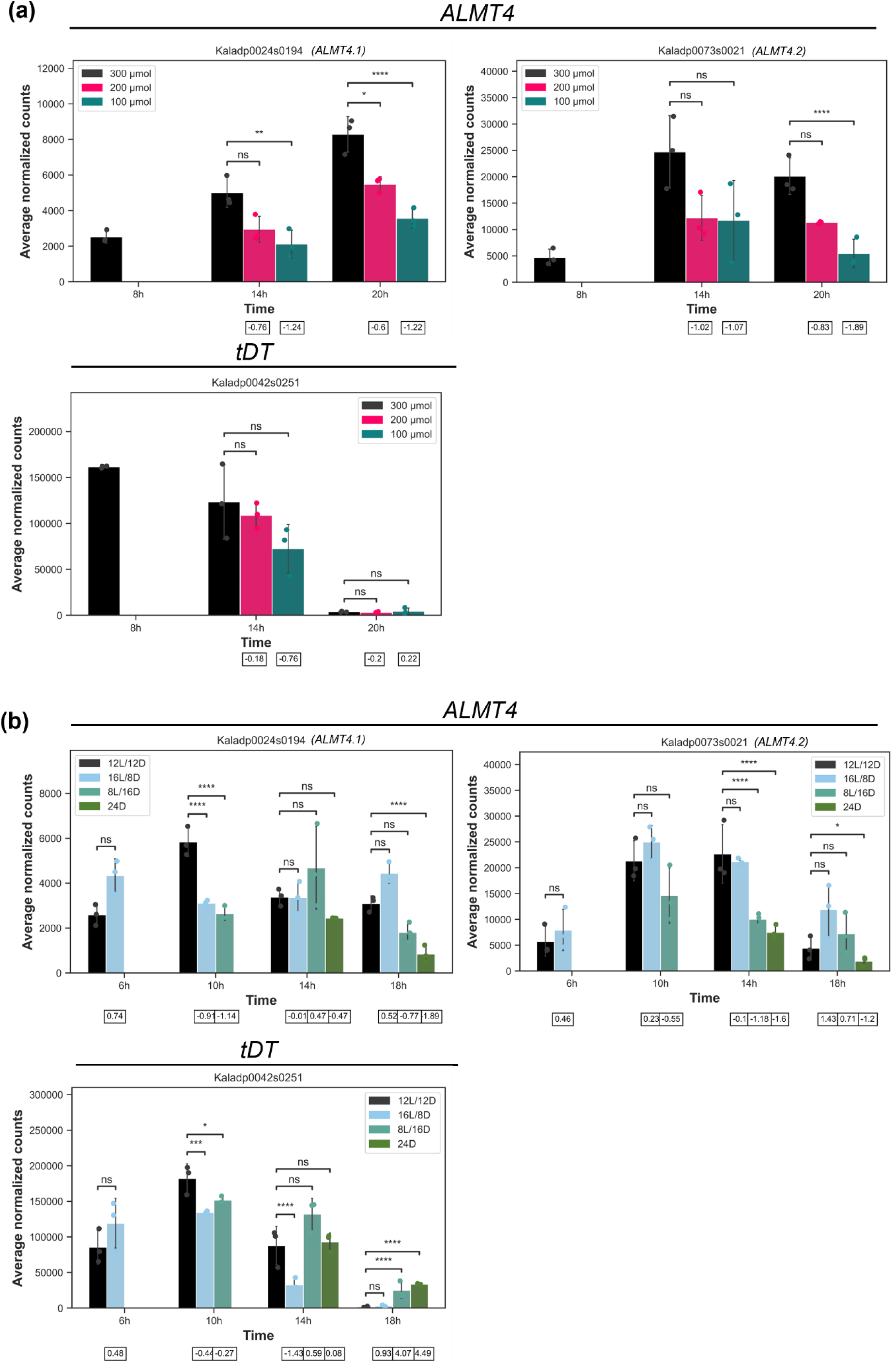
Temporal transcript abundance patterns of *ALMT4* and *tDT* genes in leaves of *Kalanchoë fedtschenkoi* under different light intensities and photoperiods. Temporal transcript abundances (average normalized counts) of *ALMT4* and *tDT* for LP5, 6, 7 of *K. fedtschenkoi* under different light intensities (300-200-100 µmol m^-2^ s^-1^) at 08:00, 14:00 and 20:00 h (a) and photoperiods at 06:00, 10:00, 14:00, 18:00 h (b). Values inside boxes below the bars represent the log2 fold change (log_2_FC) of the treatment compared to the control (300 µmol m^-2^ s^-1^ in the light intensity experiment and 12L/12D in the photoperiod experiment) (n=3 plants). Error bars represent SD. Significant differences were tested using DESeq2 (padj values) with the significance threshold P=0.05 and are indicated with an asterisk (*P<0.05; **P<0.01; ***P<0.001; ****P<0.0001). RNA-seq samples were collected at 7:45 h in darkness, since lights were switched on at 08:00 h. At this time, plants from the other treatments were also still exposed to darkness and thus received the same treatment. Therefore, data is only available for the control treatment (300 µmol m^-2^ s^-1^) at 08:00 h. The same applies to the 06:00 h and 10:00 h time points in the photoperiod experiment. At 06:00 h, transcript levels for plants under 8L/16D and 24D are expected to match those of 12L/12D, as all were exposed to darkness. Similarly, at 10:00 h, transcript levels for plants under 24D are expected to be the same as those under 8L/16D, as these plants were still all exposed to darkness.

In contrast to *ALMT* genes, *tDT* transcripts (Kaladp0042s0251, tDT is encoded by a single gene in all plant genomes sequenced to date, Wai et al., 2017) peaked early at dawn [between 08:00 (already before the light period) and 10:00 h] and then gradually declined throughout the light period, reaching near-zero levels by dusk under control conditions (300 µmol m^-2^ s^-1^ and 12L/12D) (Figs 6a, b). Under a shortened photoperiod and continuous darkness, *tDT* transcripts still peaked around similar levels at 10:00 h and showed similar abundance at 14:00 h compared to control conditions. However, under these light-limited conditions transcript levels showed a significant, massive increase around dusk (18:00 h) compared to controls (Figs 5b, 6b). In contrast, *tDT* transcripts were downregulated at 14:00 h under an extended photoperiod (16L/8D).

#### Cytosol

The *NADP-ME* gene family comprises five members in *K. fedtschenkoi*. The cytosolic forms of NADP-ME, encoded by *NADP-ME3.1* (Kaladp0102s0114) and *NADP-ME3.2* (Kaladp0024s0016), showed a diurnal transcript abundance pattern with peak abundance occurring at dawn and in the second half of the light period, respectively (Fig. S2 a, c). Both *NADP-ME* genes also exhibited sensitivity to light, with lower transcript levels at 14:00 h for *NADP-ME3.2* and generally higher abundance at each timepoint for *NADP-ME3.1* under a shortened photoperiod (8L/16D) and continuous darkness (Fig. 5, Fig. S2 c).

#### Mitochondrion

The *NAD-ME* gene family in *K. fedtschenkoi* consists of seven members, all of which maintained rather stable transcript levels throughout the light period (Fig. S3). Three *NAD-ME* gene family members with the highest transcript abundance (with *NAD-ME1.1* (Kaladp0033s0124) exhibiting the highest transcript levels), encoding mitochondrial malic enzyme subunits (NAD-ME functions as a homo-or heterodimer), also exhibited only minor changes in transcript levels under different light treatments compared to control conditions (Fig. 5).

#### Chloroplast

A gene encoding a chloroplastic form of NADP-ME, namely *NADP-ME4* (Kaladp0092s0166), showed remarkably higher transcript abundance than other *NADP-ME* gene family members and exhibited strong sensitivity to both timepoint and light conditions. In control conditions, *NADP-ME4* transcript abundance peaked at dawn and dropped to near-zero levels by dusk (Figs S2 a, c). However, under reduced light intensities, a shortened photoperiod and continuous darkness, transcript levels massively increased (log_2_FC between 5-7) in the afternoon and at dusk (Fig. 5, Fig. S2 a, c).

The *Ribulose bisphosphate carboxylase* gene family includes six members in *K. fedtschenkoi*. Most genes encoding the Rubisco large chain *(rbcL*) and small subunit 3B (*RBCS-3B*) showed relatively stable temporal transcript abundance patterns during the light period, without notable peaks at specific timepoints (Fig. S4). In addition, their transcript levels were generally unaffected by light intensity. An exception was *RBCS3B.2* (Kaladp0442s0049, encoding a small subunit), which showed a midday dip in transcript levels and strong downregulation at dusk under light-limiting conditions (8L/16D and 24D) (Fig. 5b, Fig. S4 c).

All three *RCA* genes (three *RCA* gene family members in *K. fedtschenkoi*) displayed a similar diurnal transcript abundance pattern under control light conditions, with peak abundance detected at dawn followed by a gradual decline throughout the light period (Fig. S5). These genes also exhibited light-responsive expression, with *RCA1.1* (Kaladp0033s0048) generally showing lower transcript abundance under reduced light intensities and exhibiting stable transcript levels among the different photoperiods (Fig. 5). *RCA2* (Kaladp0075s0059) and *RCA1.2* (Kaladp0083s0067) displayed upregulation particularly at 10:00, 14:00 and/or 18:00 h under a curtailed photoperiod and continuous darkness (Fig. 5, Fig. S5).

Both *PPDK* genes (two members in *K. fedtschenkoi*), *PPDK.2* (Kaladp0076s0229) and *PPDK.6* (Kaladp0039s0092), showed a comparable pattern of diurnal transcript levels and behavior to changing light conditions. Peak abundance was noticed at dawn, followed by a gradual decrease toward dusk under control conditions. Transcript levels were obviously higher throughout the light period under a shortened photoperiod and continuous darkness (Fig. 5, Figs S6 a, b).

Our RNA-seq data indicated that *K. fedtschenkoi* possesses two *PPDK-RP* genes associated with CAM. *PPDK-RP1.2* (Kaladp0060s0363) demonstrated much higher sensitivity to light and temporal regulation of its transcript levels compared to *PPDK-RP1.1* (Kaladp0010s0106) (Fig. 5, Figs S6 c, d). Both are orthologs to At4g21210, which encodes a PPDK-RP that has both protein kinase and protein phosphatase activities towards PPDK (Astley et al., 2011). A peak in *PPDK-RP1.2* transcript abundance was observed at dusk under control conditions, whereas its transcript levels remained lower throughout the light period under a shortened photoperiod and continuous darkness. Conversely, *PPDK-RP1.1* exhibited higher transcript levels than *PPDK-RP1.2* but displayed a rather stable temporal pattern of transcript abundance that remained rather unaffected by light availability or timepoint (Fig. 5, Figs S6 c, d).

## Discussion

The discussion is structured around different biochemical aspects related to the fate of malate in CAM mesophyll cells during the light period: (1) its release from the vacuole via either ALMT and/or tDT, (2) malate decarboxylation by ME, (3) refixation of CO_2_ by Rubisco and recycling of ME-derived pyruvate to starch via PPDK. By exploring the effects of light intensity and photoperiod on these biochemical events through an integrative analysis spanning physiological, metabolic, protein, enzymatic, and gene transcript levels, we aim to expand our knowledge about diurnal malate metabolism in CAM. Under control conditions, the transcript abundance patterns of our genes-of-interest (related to vacuolar malate remobilisation) proved to be consistent with those previously measured by RNA-seq and/or qRT-PCR in *K. fedtschenkoi* (Dever et al., 2015; Yang et al., 2017) and other (facultative) CAM species such as pineapple (Wai et al., 2017), *Portulaca oleracea* (Ferrari et al., 2020), and *Portulaca amilis* (Gilman et al., 2022).

### Release of malate out of the vacuole via either ALMT and/or tDT (1)

Orthologs to *AtALMT4* (At1g25480) (i.e. *KfALMT4.1* and *KfALMT4.2*) were the two most highly expressed members of the *ALMT* gene family in *K. fedtschenkoi* during daytime (Figs 6, S1). Electrophysiological approaches in Arabidopsis have confirmed that *At*ALMT4 can mediate vacuolar malate efflux in guard cells during ABA-induced stomatal closure (Eisenach et al., 2017). Since RNA extraction and sequencing were performed on bulk leaf samples in our study, it can be assumed that these genes encode tonoplastic ALMT4 in mesophyll cells rather than in guard cells. Lefoulon et al. (2020) discovered that transcript levels of *KfALMT12*, a specific member of the *ALMT* gene family, exhibit a diurnal cycle that roughly follows the temporal pattern of anion current activity in *K. fedtschenkoi* guard cells. It was recently also demonstrated that tDT protein levels matched tDT transcript levels, which were both higher during the middle of the day and reduced at midnight in *K. laxiflora* (Schiller et al., 2024). During cold acclimation in Arabidopsis, a higher malate content has been associated with an increased abundance of tDT protein (Schulze et al., 2012). All these observations indicate that *ALMT* and *tDT* transcript levels could serve as a proxy for malate transport capacity.

In our study, malate degradation was generally initiated between 10:00 and 12:00 h under control light conditions (Figs 2a, c). *KfALMT4* transcripts peaked during a similar part of the day (10:00-14:00 h), whilst *KftDT* transcripts peaked earlier around dawn (08:00-10:00 h) and reached near-zero levels (gene off) at dusk (Figs 6a, b). Based solely on these temporal transcript levels of both transporters, it remains challenging to ultimately designate the vacuolar malate exporter. Also considering a general 4-hour time shift between transcript and functional protein due to mRNA processing and translation (as also observed by Hartwell et al., 1996, 1999 for *KfPPCK1* and suggested by Lefoulon et al., 2020 for *KfALMT12*), both transporters remain likely candidates. However, several observations in our paper point to ALMT4 as a more plausible candidate mediating vacuolar malate efflux in CAM. (i) Premature malate degradation was not observed between 04:00 and 08:00 h when the photoperiod was extended by switching on the lights sooner at 04:00 h (16L/8D) (Fig. 2c). *KfALMT4* transcript levels were neither significantly affected by this condition, whilst *tDT* transcripts clearly declined earlier under an extended photoperiod (Figs 5b, 6b). The absence of premature malate degradation in this time frame also indicates that the activity of the tonoplast malate exporter, or a downstream actor such as ME, is rapidly activated around the expected dawn (i.e. 8:00 h), probably via a circadian clock-associated post-translational modification (PTM). In the C_3_ plant Arabidopsis, *At*ALMT4 has been reported to possess a specific C-terminal serine (S382) that can be phosphorylated (inactivation of channel activity) by mitogen-activated protein kinases (MAPKs) *in vitro* (Eisenach et al., 2017). Also for *At*tDT a specific phosphorylation site has already been reported (S409) (Schulze et al., 2012), but the impact of its (de)phosphorylation on activity remains yet to be confirmed. (ii) Shortening the photoperiod by providing illumination only at 12:00 h (8L/16D) caused a consistent delay in malate degradation but plants were still able to degrade similar amounts of malate in the early afternoon, followed by a similar Phase IV CO_2_ uptake pattern in comparison to controls (Figs 1b, 2c). Under these conditions, *KfALMT4* genes were also significantly downregulated during the morning period, whilst *tDT* transcript abundance was not influenced during the major part of the day (Figs 5b, 6b). (iii) While transcript levels of both *KfALMT4* genes were significantly downregulated under the reduced 100 µmol m^-2^ s^-1^, *tDT* transcript abundance remained unaffected (Figs 5a, 6a). This reduction in light intensity also had a notable impact on CO_2_ uptake, malate degradation and starch accumulation, and the activity levels of NAD-ME, Rubisco, and PPDK (Figs 1a, 2a, b, 3a, c, d). Taken together, these integrated findings (i, ii, iii) clearly demonstrate a closer correlation between *ALMT4* transcript patterns and observations at the physiological and biochemical level compared to *tDT* mRNA patterns. This suggests that ALMT4 plays a more prominent role in diurnal CAM than tDT, making it a more reasonable candidate to mediate vacuolar malate efflux. Furthermore, our experiments indicate that vacuolar malate remobilisation is primarily regulated by the endogenous clock and photoperiod, with light intensity playing an additional role.

### Decarboxylation of malate by ME (2)

All *NAD-ME* genes displayed relatively stable transcript levels throughout the light period and were rather insensitive to the different light conditions, indicating minor temporal- and light-dependent transcriptional regulation (Figs 5, S3). Mitochondrial NAD-ME was the primary decarboxylase, exhibiting ca. five to eight times higher intrinsic activity than cytosolic/chloroplastic NADP-ME, depending on the timepoint (Figs 3a, b, e, f). The strong light-dependent and temporal regulation of chloroplastic *KfNADP-ME4* and cytosolic *KfNADP-ME3.1* at the transcript level was not mirrored in the measured enzyme activities, which were largely insufficient to accommodate the observed malate decarboxylation. These observations align with previous reported activities of NAD-ME and NADP-ME in the CAM species *K. fedtschenkoi* and *Phalaenopsis* ‘Edessa’ (Dever et al., 2015; Daems et al., 2024). In our study, NAD-ME activity increased sooner under an extended photoperiod and later under a shortened shortened one (Fig. 3e), illustrating the significant influence of photoperiod on malate decarboxylation, with light intensity playing an additional role. NAD-ME activities were also found to be in excess compared to the observed malate degradation rates. For example, under control light conditions, the maximum malate decarboxylation rate was 25 ± 7 µmol g^-1^FW malate/2h around 12:00 h (Fig. 2c), whereas NAD-ME activities were 14 ± 0.8 µmol PYR h^-1^ g^-1^FW (equivalent to 28 µmol PYR 2h^-1^ g^-1^FW) (Fig. 3e). Even at low light intensities (100, 50, and 10 µmol m^-2^ s^-1^) and continuous darkness, NAD-ME activity remained relatively high (ca. 12 µmol PYR h^-1^ g^-1^FW), implying that *in vitro* activities of the primary decarboxylase do not impose a bottleneck for malate remobilisation.

However, decarboxylating enzyme activity might be modified *in vivo* by cellular conditions such as substrate availability, pH, energy charge and/or reversible PTMs to exert its activity exclusively during the day. Numerous PTMs have been identified for mitochondrial NAD-ME and cytosolic NADP-ME in Arabidopsis, though their functional roles remain unvalidated (Schiller and Bräutigam, 2021). In C_4_ plants, plastidic NADP-ME is redox-regulated and active in its reduced form (Drincovich and Andreo, 1994; Alvarez et al., 2012). The cysteine residues responsible for this redox regulation are conserved in plastidic NADP-MEs. Activation of NADP-ME is thus connected to a reduced plastid stroma and influenced by light intensity through photosynthetic reduction of the plastid stroma. It remains to be established whether a similar redox regulation mechanism applies to mitochondrial NAD-ME in CAM, where a reduced mitochondrial matrix could activate NAD-ME during the light period.

### Refixation of CO_2_ by Rubisco and recycling of ME-derived pyruvate to starch via PPDK (3)

#### CO_2_ assimilation by Rubisco

Under control conditions, intrinsic Rubisco activity was ca. 6 µmol CO_2_ h^-1^ g^-1^FW at dawn and increased to maximal levels (ca. 14 µmol CO_2_ h^-1^ g^-1^FW) during the latter half of the light period (Figs 3c, g; consistent with observations by Maxwell et al., 1999; Maxwell et al., 2002). These measurements indicate that Rubisco activity seemed always sufficient to capture the ME-liberated CO_2_ (maximal malate degradation at noon: 25 ± 7 µmol g^-1^FW malate 2h^-1^ versus Rubisco activity of 10.5 ± 2.5 µmol CO_2_ h^-1^ g^-1^FW) (Figs 2a, c). Rubisco activity was also clearly affected by both light intensity and photoperiod. Under light-limiting conditions (100, 50, 10 µmol m^-2^ s^-1^ and continuous darkness) the expected stimulation of Rubisco activity during the day did not occur (Figs 3c, g). Though not observed, these activity levels were still sufficiently high to allow for some degree of malate decarboxylation, suggesting that intrinsic Rubisco activity could not have limited malate remobilisation under such conditions.

The substantial rise in Phase II carbon fixation between 04:00 and 07:30 h under an extended photoperiod (Fig. 1b) was not associated with an increase in intrinsic Rubisco activity at 06:00 h (Fig. 3g). Therefore, this increase was likely driven by a light-induced decrease in the stromal ADP/ATP ratio, which can activate Rubisco via increased Rubisco activase activity (Zhang and Portis, 1999). Under this photoperiodic condition, total Phase II CO_2_ uptake rate (04:00-07:30 h) was 67.5 ± 12.5 mmol CO_2_ m^-2^, with PEPC and Rubisco contributing 32.1 ± 3.6 and ca. 35 mmol CO_2_ m^-2^, respectively (Fig. 1b). At 06:00 h, Rubisco activity reached 5.7 ± 1.3 µmol CO_2_ h^-1^ g^-1^FW (equivalent to 14.1 ± 3.2 mmol CO_2_ h^-1^ m^-2^ or 49.4 mmol CO_2_ 3.5h^-1^ m^-2^). This base level of intrinsic Rubisco activity was clearly sufficient to accommodate the increased CO_2_ fixation rate. This earlier onset of Rubisco-mediated CO_2_ uptake was also mirrored in significantly higher starch and sucrose levels at 08:00 h compared to controls (Figs 2d, e). Remarkably, plants under both an extended and shortened photoperiod showed a similar amount of CO_2_ uptake during Phase IV compared to controls (Fig. 1b), which was also reflected in similar Rubisco activities among these treatments at 16:00 and 20:00 h (Fig. 3g). This demonstrates that light intensity was more important than photoperiod to mediate Phase IV carbon fixation.

A notably similar temporal and light-responsive intrinsic activity pattern to that of Rubisco (CO_2_ fixation) was observed for mitochondrial NAD-ME (CO_2_ liberation), with both enzymes exhibiting a doubling in activity during the latter part of the light phase under control light conditions (Figs 3a, c, e, g). This indicates the possibility of coordinated regulation via an as-yet unidentified shared mechanism in CAM. Redox regulation of both enzymes (NAD-ME in the matrix, Rubisco in the stroma) is a strong candidate, particularly given the well-established redox regulation of Rubisco in Arabidopsis (Zhang et al., 2002) and plastidial NADP-ME in C_4_ plants (Drincovich and Andreo, 1994; Alvarez et al., 2012).

### Recycling of ME-derived pyruvate to starch via PPDK

Under control light conditions, PPDK activity was minimal at dawn (ca. 1 µmol PEP h^-1^ g^-1^FW) but exhibited a massive increase to 7.4 ± 1.3 µmol PEP h^-1^ g^-1^FW (equivalent to 14.8 µmol PEP 2h^-1^ g^-1^FW) between 10:00 and 12:00 h, which persisted until 16:00 h (Figs 3d, h). This pattern closely mirrored the dynamics of malate, with maximum degradation occurring around 12:00 h (25 ± 7 µmol g^-1^FW malate 2h^-1^) (Figs 2a, c). This elevated PPDK activity also supported high rates of starch accumulation during this part of the photoperiod (Figs 2b, d). At lower light intensities (50 and 10 µmol m^-2^ s^-1^) and in 24D, PPDK activity and malate content remained notably low and high respectively throughout the light period, similar to their original dawn levels (Figs 2a, c, 3d, h). Under these low light conditions, also starch and sucrose concentrations remained low (Figs 2b, d, e). Under an extended photoperiod, PPDK activity massively increased between 8:00 and 12:00 h (to ca. 10.2 ± 2.4 µmol PEP h^-1^ g^-1^FW) (Fig. 3h), while the highest malate degradation rate was also observed within this time frame (19.8 ± 3.1 µmol malate g^-1^FW 2h^-1^) (Fig. 2c). Under a shortened photoperiod, PPDK activity exceeded the maximum levels observed under control conditions during the final part of the photoperiod (13.3 ± 2.7 µmol PEP h^-1^ g^-1^FW versus 8.6 ± 1.9 µmol PEP h^-1^ g^-1^FW at 16:00 h) (Fig. 3h), with malate degradation also peaking during this time frame (17.1 ± 4.2 µmol malate g^-1^FW 2h^-1^) (Fig. 2c). In addition, the endogenous clock seemed also to play a role in regulating PPDK activity, since no large premature increase was noticed between 4:00 and 8:00 h (Fig. 3h), consistent with the absence of premature malate degradation (Fig. 2c). In summary, these findings uncover a strikingly close light-sensitive correlation between the dynamics of malate and PPDK activity levels, suggesting that this recycling enzyme could act as a potential limiting factor for malate remobilisation, probably via a feedback mechanism. Under light-limited conditions, ME-derived pyruvate may accumulate due to a diminished PPDK activity, leading to feedback inhibition of ME and consequently also hindering further vacuolar malate efflux. These findings strengthen and refine a previously mentioned hypothesis that the extractable activities of PPDK might be sufficiently low to potentially be at least partially rate-limiting in the overall deacidification process (Holtum and Osmond, 1981; Neuhaus et al., 1988; Holtum et al., 2005).

The pronounced light and temporal sensitivity of PPDK abundances and activities (Figs 3d, h, 4) also underscores a major role of its upstream regulator, PPDK-RP. Despite *KfPPDK-RP1.1* exhibited the highest transcript levels of both *PPDK-RP* genes in *K. fedtschenkoi*, we propose only a minor contribution to PPDK regulation given its relatively stable transcript levels across all time points and insensitivity to the applied light conditions (Figs 5, S6c, d). In contrast, *KfPPDK-RP1.2* exhibited much higher sensitivity to light and temporal regulation of its transcript levels (Figs 5b, S6c, d). Since PPDK activity and abundance were also strongly affected by light intensity, photoperiod and time point (Figs 3d, h, 4), it is highly likely that *KfPPDK-RP1.2* potentially encodes the key RP responsible for the light-dependent and temporal regulation of both cytosolic and chloroplastic PPDK in *K. fedtschenkoi*. The results demonstrated strong light-dependent regulation (both light intensity and photoperiod) of PPDK at the gene transcript, protein abundance, and intrinsic enzyme activity level, suggesting that PPDK and its RP play crucial roles in the temporal coordination of decarboxylation with malate efflux from the vacuole during the light period in CAM plants.

Taken together, the results presented here uncovered that: (1) Malate transport out of the vacuole is primarily influenced by the endogenous clock and photoperiod, with *Kf*ALMT4 being a more plausible transporter candidate than *Kf*tDT. (2) Malate decarboxylation is mainly influenced by photoperiod, with light intensity playing a supplementary role. (3) Both photoperiod and light intensity greatly affect CO_2_ refixation and pyruvate recycling, with PPDK—being the last in line—representing the most strictly light-regulated enzyme potentially governing diurnal deacidification in CAM leaves.

## Acknowledgements

Funding was provided by the Research Fund KU Leuven. Figures were created with BioRender.com.

## Author contributions

JC, BVdP, and SD designed the research; SD performed the experiments and collected the data; all authors analysed data; SD, BVdP, and JC wrote the paper; SD created the figures. All authors revised the manuscript and approved the final version.

## Data availability

The authors declare that all data supporting the findings of this study are available within the paper and its Supporting Information files (Figs S1-S6, Table S1). Sequence data from the RNA-seq experiment will be available in a public repository upon acceptance of the article.

## Supporting information

**Fig. S1** Temporal transcript abundance (average normalized counts) patterns of *ALMT* gene family members in leaves of *Kalanchoë fedtschenkoi* under different light intensities and photoperiods.

**Fig. S2** Temporal transcript abundance (average normalized counts) patterns of *NADP-ME* gene family members in leaves of *Kalanchoë fedtschenkoi* under different light intensities and photoperiods.

**Fig. S3** Temporal transcript abundance (average normalized counts) patterns of *NAD-ME* gene family members in leaves of *Kalanchoë fedtschenkoi* under different light intensities and photoperiods.

**Fig. S4** Temporal transcript abundance (average normalized counts) patterns of *Rubisco* in leaves of *Kalanchoë fedtschenkoi* under different light intensities and photoperiods.

**Fig. S5** Temporal transcript abundance (average normalized counts) patterns of *Rubisco activase (RCA)* in leaves of *Kalanchoë fedtschenkoi* under different light intensities and photoperiods.

**Fig. S6** Temporal transcript abundance (average normalized counts) patterns of *PPDK and PPDK-RP* in leaves of *Kalanchoë fedtschenkoi* under different light intensities and photoperiods.

**Table S1** Slope coefficients as a measure for the rates of malate degradation and starch accumulation.

**Methods S1** Extended Materials and Methods.

