## Supporting Information for "The last one in the row but decisive: PPDK as a potential key regulator of diurnal deacidification in CAM leaves across varying PPFD and photoperiod conditions"

The following Supporting Information is available for this article:

**Fig. S1** Temporal transcript abundance (average normalized counts) patterns of *ALMT* gene family members in leaves of *Kalanchoë fedtschenkoi* under different light intensities and photoperiods.

**Fig. S2** Temporal transcript abundance (average normalized counts) patterns of *NADP-ME* gene family members in leaves of *Kalanchoë fedtschenkoi* under different light intensities and photoperiods.

**Fig. S3** Temporal transcript abundance (average normalized counts) patterns of *NAD-ME* gene family members in leaves of *Kalanchoë fedtschenkoi* under different light intensities and photoperiods.

**Fig. S4** Temporal transcript abundance (average normalized counts) patterns of *Rubisco* in leaves of *Kalanchoë fedtschenkoi* under different light intensities and photoperiods.

**Fig. S5** Temporal transcript abundance (average normalized counts) patterns of *Rubisco activase (RCA)* in leaves of *Kalanchoë fedtschenkoi* under different light intensities and photoperiods.

**Fig. S6** Temporal transcript abundance (average normalized counts) patterns of *PPDK and PPDK-RP* in leaves of *Kalanchoë fedtschenkoi* under different light intensities and photoperiods.

**Table S1** Slope coefficients as a measure for the rates of malate degradation and starch accumulation.

**Methods S1** Extended Materials and Methods.

**Fig. S1 Temporal transcript abundance (average normalized counts) patterns of *ALMT* gene family members in leaves of *Kalanchoë fedtschenkoi* under different light intensities and photoperiods.** *ALMT* transcript levels for LP5, 6, 7 of *K. fedtschenkoi* under different light intensities (300-200-100 µmol $m^{-2}$ $s^{-1}$) at 08:00, 14:00 and 20:00 h (a) and photoperiods at 06:00, 10:00, 14:00, 18:00 h (b). Both panels (a) and (b) depict transcript abundance patterns of gene family members that were not included in the heatmaps (Fig. 5). Values inside boxes below the bars represent the log2 fold change ($log_2FC$) of the treatment compared to the control (300 µmol $m^{-2}$ $s^{-1}$ in the light intensity experiment and 12L/12D in the photoperiod experiment) (n=3 plants). Error bars represent SD. Significant differences were tested using DESeq2 (padj values) with the significance threshold P=0.05 and are indicated with an asterisk (*P<0.05; **P<0.01; ***P<0.001; ****P<0.0001). RNA-seq samples were collected at 7:45 h in darkness, since lights were switched on at 08:00 h. At this time, plants from the other treatments were also still exposed to darkness and thus received the same treatment. Therefore, data is only available for the control treatment (300 µmol $m^{-2}$ $s^{-1}$) at 08:00 h. The same applies to the 06:00 h and 10:00 h time points in the photoperiod experiment. At 06:00 h, transcript levels for plants under 8L/16D and 24D are expected to match those of 12L/12D, as all were exposed to darkness. Similarly, at 10:00 h, transcript levels for plants under 8L/16D are expected to be same as those under 24D, as these plants were still all exposed to darkness.

*ALMT*

**(a)**

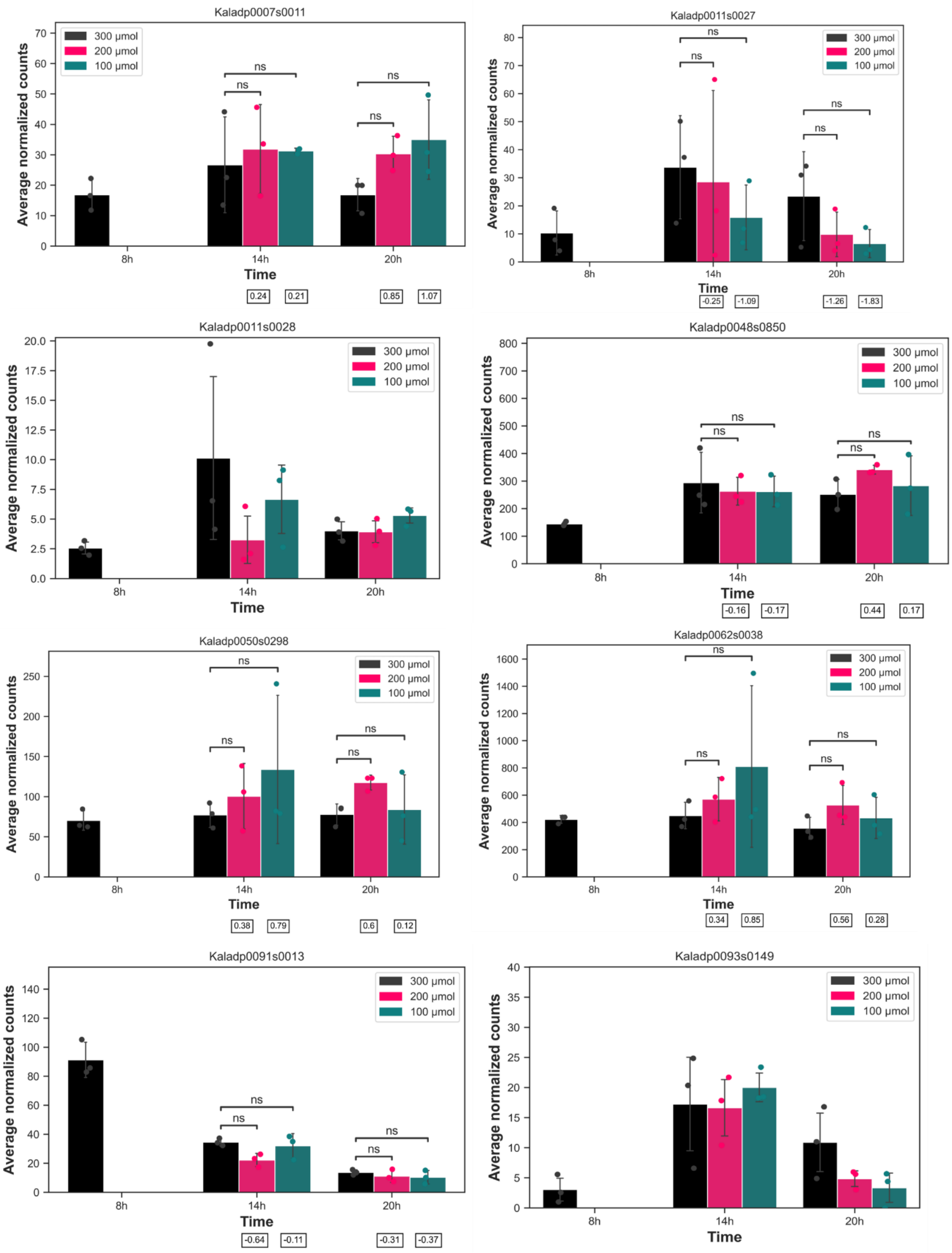

Kaladp0007s0011
300 μmol
200 μmol
100 μmol
Average normalized counts
ns
8h
14h
20h
Time
0.24
0.21
0.85
1.07
Kaladp0011s0027
-0.25
-1.09
-1.26
-1.83
Kaladp0011s0028
Kaladp0048s0850
-0.16
-0.17
0.44
0.17
Kaladp0050s0298
0.38
0.79
0.6
0.12
Kaladp0062s0038
0.34
0.85
0.56
0.28
Kaladp0091s0013
-0.64
-0.11
-0.31
-0.37
Kaladp0093s0149

**(b)**

*ALMT*

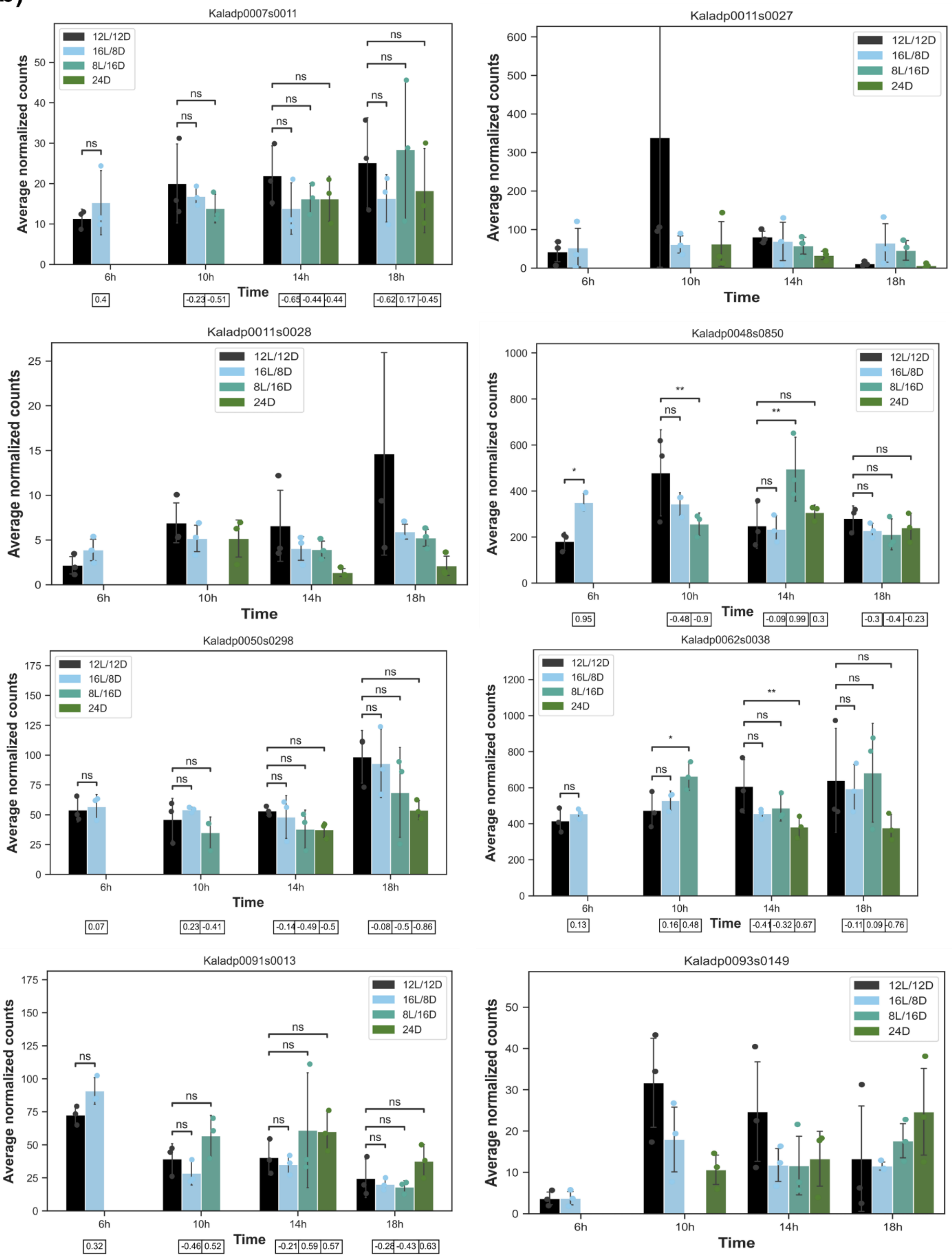
Kaladp0007s0011
Kaladp0011s0027
Kaladp0011s0028
Kaladp0048s0850
Kaladp0050s0298
Kaladp0062s0038
Kaladp0091s0013
Kaladp0093s0149
12L/12D
16L/8D
8L/16D
24D
Average normalized counts
Time
6h
10h
14h
18h
ns
*
**
0.4
-0.23 -0.51
-0.65 -0.44 -0.44
-0.62 0.17 -0.45
0.95
-0.48 -0.9
-0.09 0.99 0.3
-0.3 -0.4 -0.23
0.07
0.23 -0.41
-0.14 -0.49 -0.5
-0.08 -0.5 -0.86
0.13
0.16 0.48
-0.41 -0.32 0.67
-0.11 0.09 -0.76
0.32
-0.46 0.52
-0.21 0.59 0.57
-0.28 -0.43 0.63

**Fig. S2 Temporal transcript abundance (average normalized counts) patterns of *NADP-ME* gene family members in leaves of *Kalanchoë fedtschenkoi* under different light intensities and photoperiods.** *NADP-ME* transcript levels for LP5, 6, 7 of *K. fedtschenkoi* under different light intensities (300-200-100 µmol $m^{-2}$ $s^{-1}$) at 08:00, 14:00 and 20:00 h (a, b) and photoperiods at 06:00, 10:00, 14:00, 18:00 h (c, d). Panels (a) and (c) depict transcript abundance patterns of gene family members included in the heatmaps (Fig. 5). Values inside boxes below the bars represent the log2 fold change ($log_2FC$) of the treatment compared to the control (300 µmol $m^{-2}$ $s^{-1}$ in the light intensity experiment and 12L/12D in the photoperiod experiment) (n=3 plants). Error bars represent SD. Significant differences were tested using DESeq2 (padj values) with the significance threshold P=0.05 and are indicated with an asterisk (*P<0.05; **P<0.01; ***P<0.001; ****P<0.0001). RNA-seq samples were collected at 7:45 h in darkness, since lights were switched on at 08:00 h. At this time, plants from the other treatments were also still exposed to darkness and thus received the same treatment. Therefore, data is only available for the control treatment (300 µmol $m^{-2}$ $s^{-1}$) at 08:00 h. The same applies to the 06:00 h and 10:00 h time points in the photoperiod experiment. At 06:00 h, transcript levels for plants under 8L/16D and 24D are expected to match those of 12L/12D, as all were exposed to darkness. Similarly, at 10:00 h, transcript levels for plants under 8L/16D are expected to be same as those under 24D, as these plants were still all exposed to darkness.

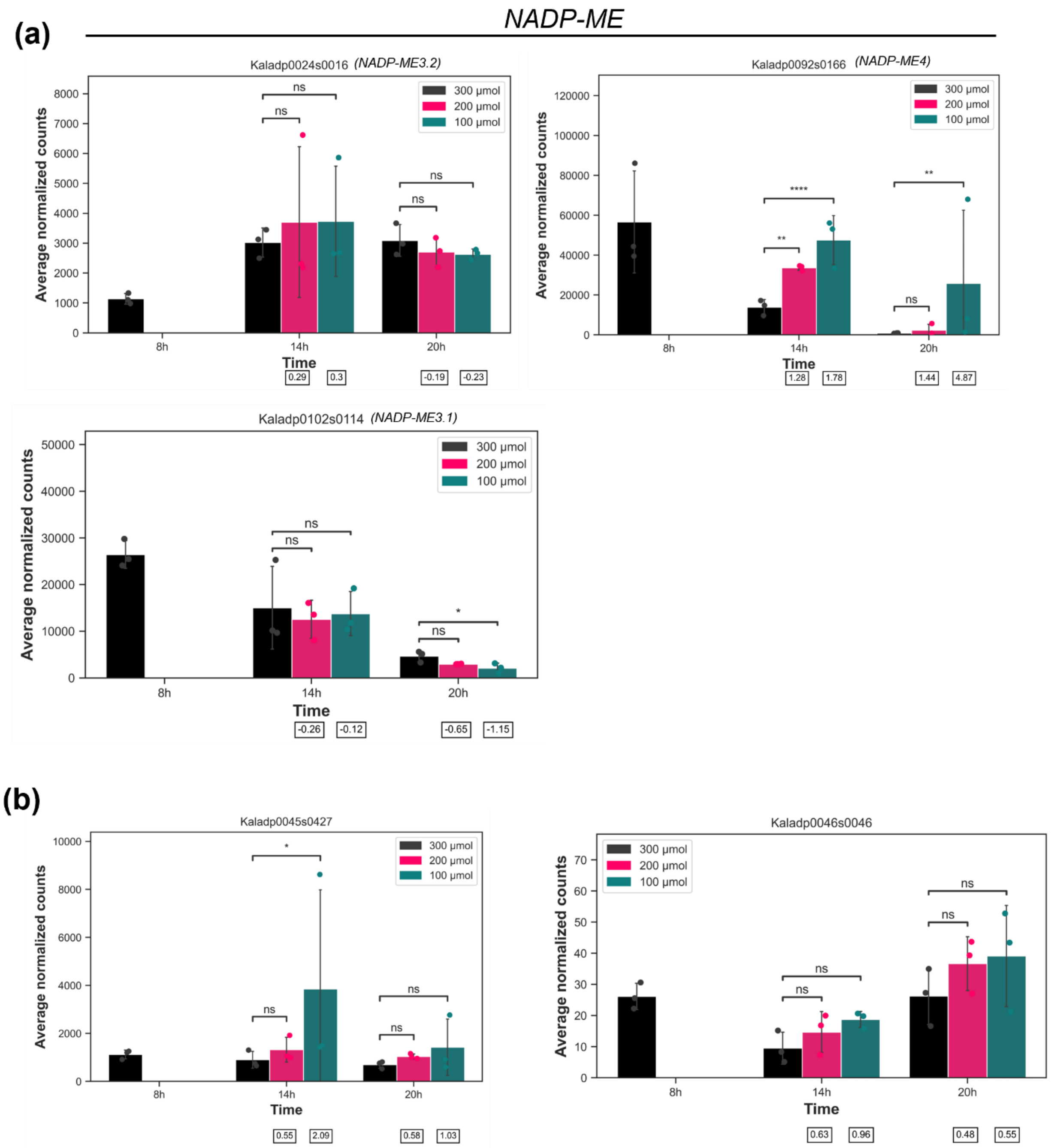

(a)
NADP-ME
Kaladp0024s0016 (NADP-ME3.2)
Average normalized counts
Time
8h
14h
20h
300 µmol
200 µmol
100 µmol
ns
0.29
0.3
-0.19
-0.23
Kaladp0092s0166 (NADP-ME4)
****
**
1.28
1.78
1.44
4.87
Kaladp0102s0114 (NADP-ME3.1)
*
-0.26
-0.12
-0.65
-1.15
(b)
Kaladp0045s0427
0.55
2.09
0.58
1.03
Kaladp0046s0046
0.63
0.96
0.48
0.55

**(c)** *NADP-ME*

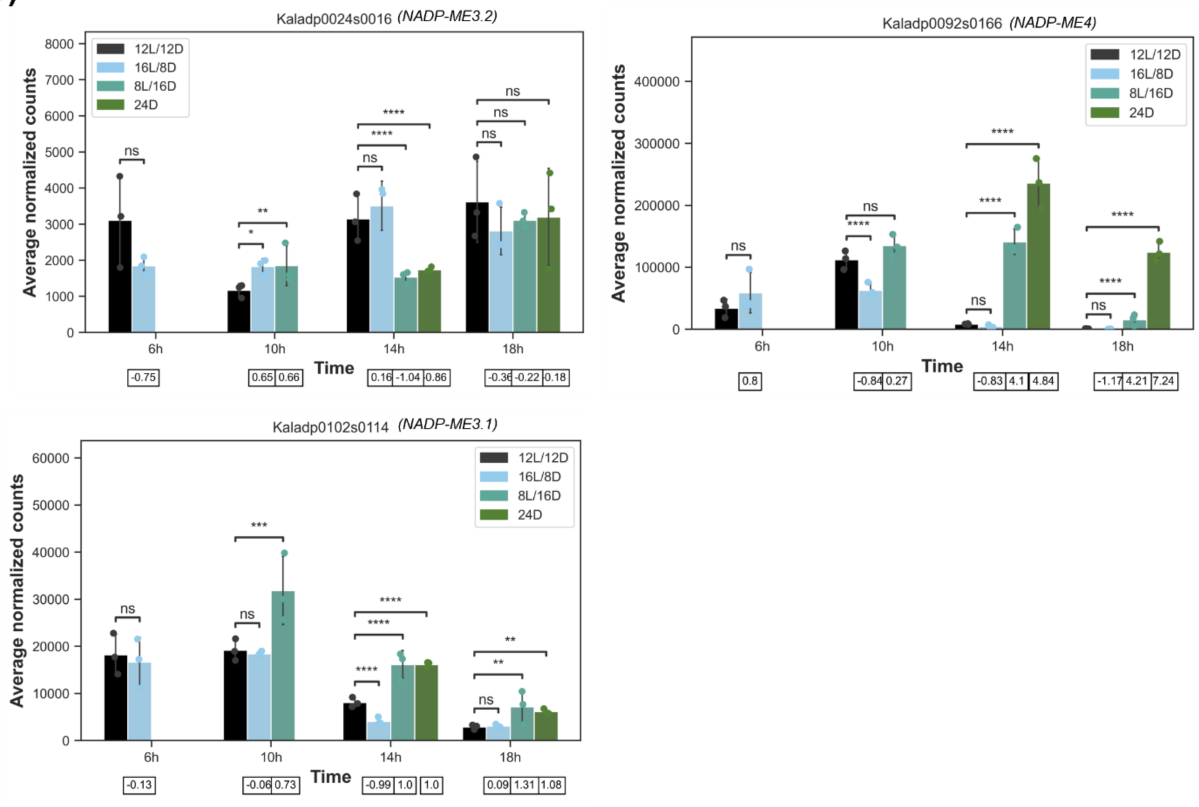
Kaladp0024s0016 (NADP-ME3.2)
Kaladp0092s0166 (NADP-ME4)
Kaladp0102s0114 (NADP-ME3.1)
12L/12D
16L/8D
8L/16D
24D
Average normalized counts
Time
6h
10h
14h
18h
-0.75
0.65 0.66
0.16 -1.04 0.86
-0.36 -0.22 0.18
0.8
-0.84 0.27
-0.83 4.1 4.84
-1.17 4.21 7.24
-0.13
-0.06 0.73
-0.99 1.0 1.0
0.09 1.31 1.08

**(d)**

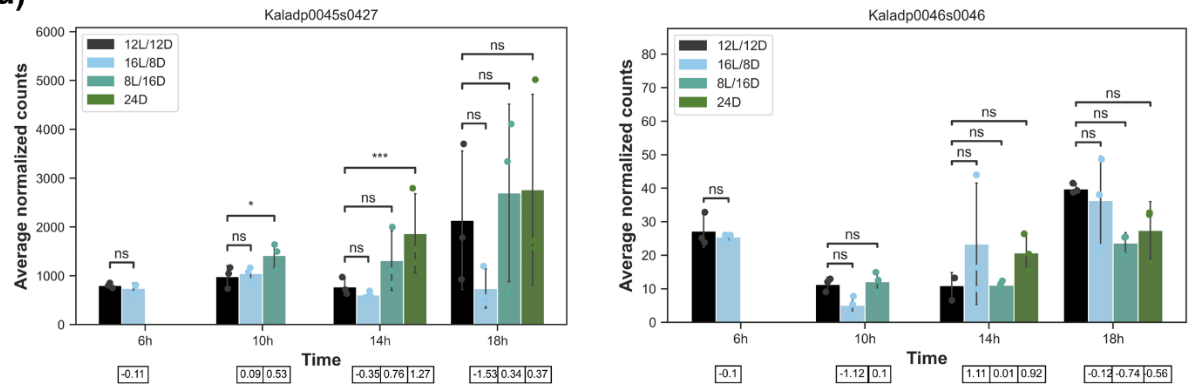
Kaladp0045s0427
Kaladp0046s0046
12L/12D
16L/8D
8L/16D
24D
Average normalized counts
Time
6h
10h
14h
18h
-0.11
0.09 0.53
-0.35 0.76 1.27
-1.53 0.34 0.37
-0.1
-1.12 0.1
1.11 0.01 0.92
-0.12 -0.74 -0.56

**Fig. S3 Temporal transcript abundance (average normalized counts) patterns of *NAD-ME* gene family members in leaves of *Kalanchoë fedtschenkoi* under different light intensities and photoperiods.** *NAD-ME* transcript levels for LP5, 6, 7 of *K. fedtschenkoi* under different light intensities (300-200-100 µmol $m^{-2}$ $s^{-1}$) at 08:00, 14:00 and 20:00 h (a, b) and photoperiods at 06:00, 10:00, 14:00, 18:00 h (c, d). Panels (a) and (c) depict transcript abundance patterns of gene family members included in the heatmaps (Fig. 5). Values inside boxes below the bars represent the log2 fold change ($log_2FC$) of the treatment compared to the control (300 µmol $m^{-2}$ $s^{-1}$ in the light intensity experiment and 12L/12D in the photoperiod experiment) (n=3 plants). Error bars represent SD. Significant differences were tested using DESeq2 (padj values) with the significance threshold P=0.05 and are indicated with an asterisk (*P<0.05; **P<0.01; ***P<0.001; ****P<0.0001). RNA-seq samples were collected at 7:45 h in darkness, since lights were switched on at 08:00 h. At this time, plants from the other treatments were also still exposed to darkness and thus received the same treatment. Therefore, data is only available for the control treatment (300 µmol $m^{-2}$ $s^{-1}$) at 08:00 h. The same applies to the 06:00 h and 10:00 h time points in the photoperiod experiment. At 06:00 h, transcript levels for plants under 8L/16D and 24D are expected to match those of 12L/12D, as all were exposed to darkness. Similarly, at 10:00 h, transcript levels for plants under 8L/16D are expected to be same as those under 24D, as these plants were still all exposed to darkness.

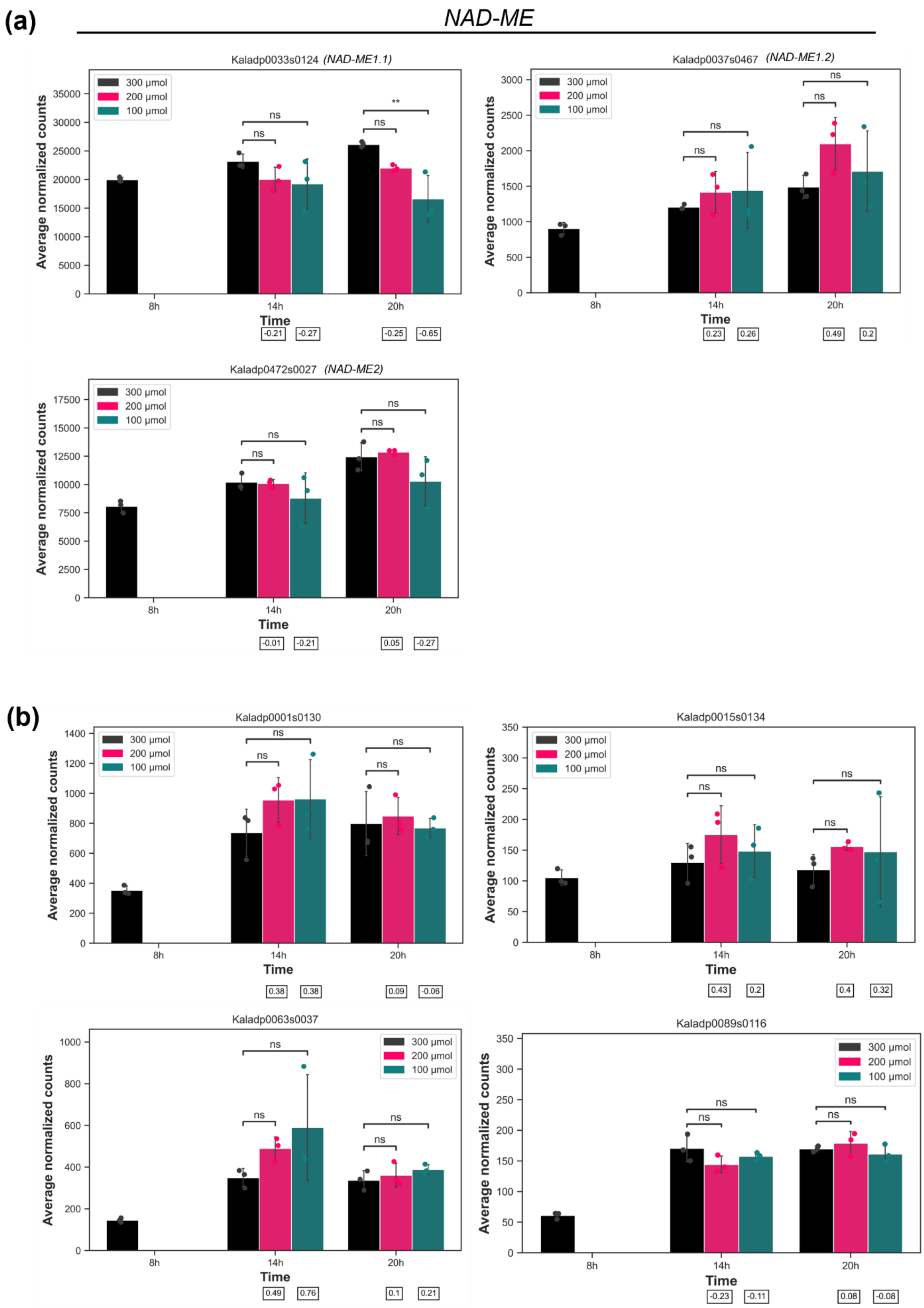
(a)
NAD-ME
Kaladp0033s0124 (NAD-ME1.1)
Kaladp0037s0467 (NAD-ME1.2)
Kaladp0472s0027 (NAD-ME2)
300 µmol
200 µmol
100 µmol
Average normalized counts
Time
8h
14h
20h
ns
**
-0.21
-0.27
-0.25
-0.65
0.23
0.26
0.49
0.2
-0.01
-0.21
0.05
-0.27
(b)
Kaladp0001s0130
Kaladp0015s0134
Kaladp0063s0037
Kaladp0089s0116
0.38
0.38
0.09
-0.06
0.43
0.2
0.4
0.32
0.49
0.76
0.1
0.21
-0.23
-0.11
0.08
-0.08

### *NAD-ME*

(c)

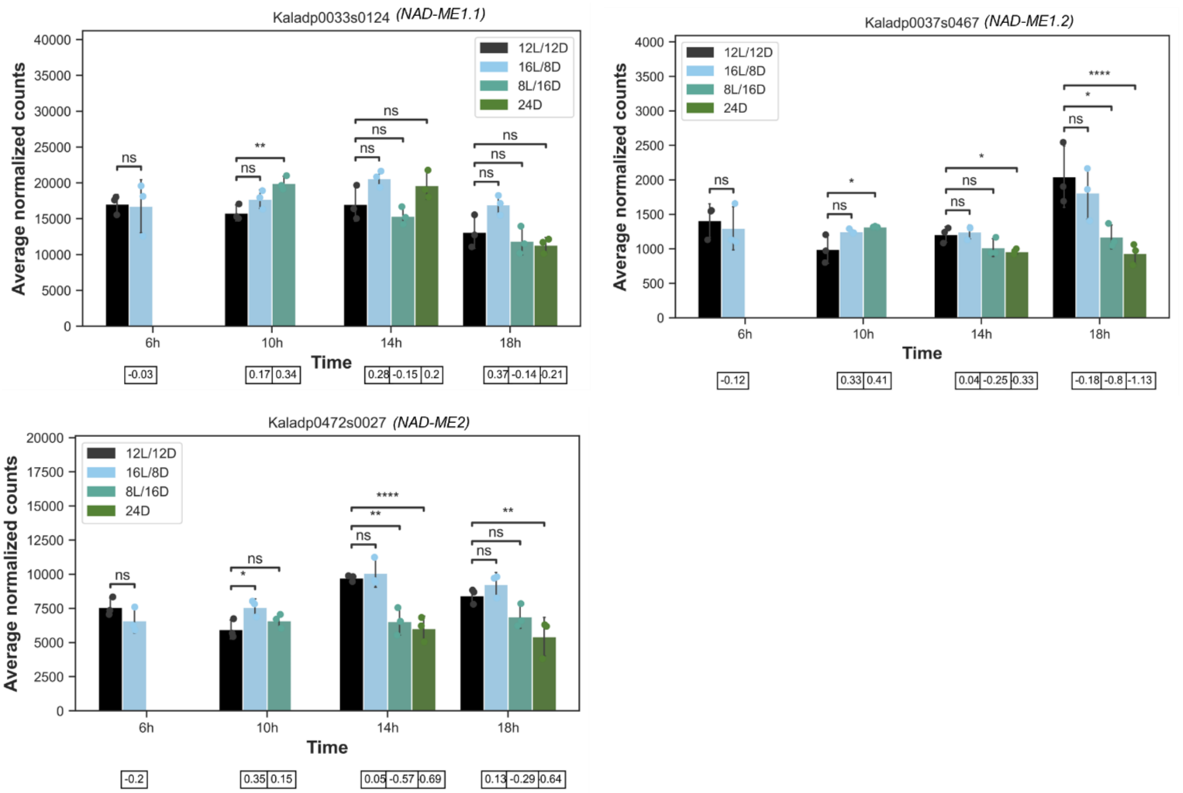
Kaladp0033s0124 (NAD-ME1.1)
Kaladp0037s0467 (NAD-ME1.2)
Kaladp0472s0027 (NAD-ME2)
Average normalized counts
Time
6h
10h
14h
18h
12L/12D
16L/8D
8L/16D
24D
ns
*
**
****
-0.03
0.17 0.34
0.28 -0.15 0.2
0.37 -0.14 0.21
-0.12
0.33 0.41
0.04 -0.25 0.33
-0.18 -0.8 -1.13
-0.2
0.35 0.15
0.05 -0.57 0.69
0.13 -0.29 0.64

(d)

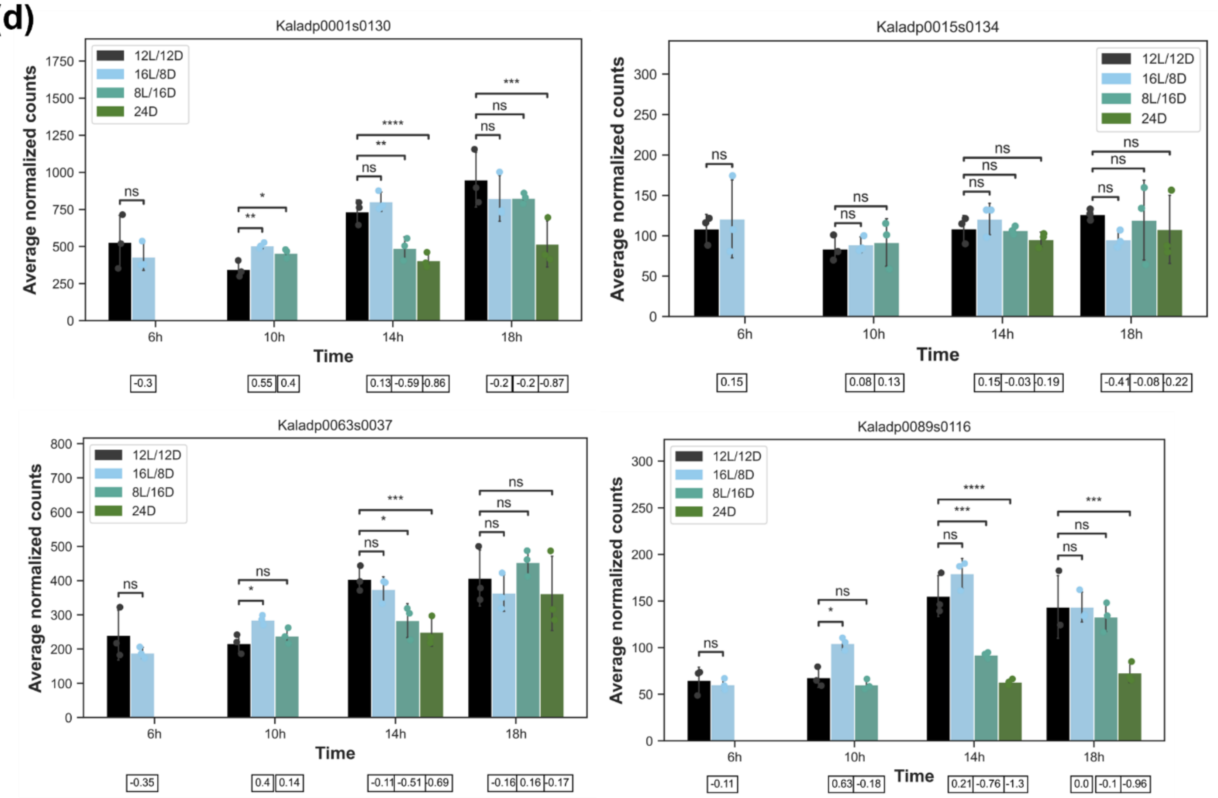
Kaladp0001s0130
Kaladp0015s0134
Kaladp0063s0037
Kaladp0089s0116
Average normalized counts
Time
6h
10h
14h
18h
12L/12D
16L/8D
8L/16D
24D
ns
*
**
***
****
-0.3
0.55 0.4
0.13 -0.59 0.86
-0.2 -0.2 -0.87
0.15
0.08 0.13
0.15 -0.03 0.19
-0.41 -0.08 0.22
-0.35
0.4 0.14
-0.11 -0.51 0.69
-0.16 0.16 -0.17
-0.11
0.63 -0.18
0.21 -0.76 -1.3
0.0 -0.1 -0.96

**Fig. S4 Temporal transcript abundance (average normalized counts) patterns of *Rubisco* in leaves of *Kalanchoë fedtschenkoi* under different light intensities and photoperiods.** *Rubisco* transcript levels for LP5, 6, 7 of *K. fedtschenkoi* under different light intensities (300-200-100 µmol $m^{-2}$ $s^{-1}$) at 08:00, 14:00 and 20:00 h (a, b) and photoperiods at 06:00, 10:00, 14:00, 18:00 h (c, d). Panels (a) and (c) depict transcript abundance patterns of gene family members included in the heatmaps (Fig. 5). Values inside boxes below the bars represent the log2 fold change ($log_2FC$) of the treatment compared to the control (300 µmol $m^{-2}$ $s^{-1}$ in the light intensity experiment and 12L/12D in the photoperiod experiment) (n=3 plants). Error bars represent SD. Significant differences were tested using DESeq2 (padj values) with the significance threshold P=0.05 and are indicated with an asterisk (*P<0.05; **P<0.01; ***P<0.001; ****P<0.0001). RNA-seq samples were collected at 7:45 h in darkness, since lights were switched on at 08:00 h. At this time, plants from the other treatments were also still exposed to darkness and thus received the same treatment. Therefore, data is only available for the control treatment (300 µmol $m^{-2}$ $s^{-1}$) at 08:00 h. The same applies to the 06:00 h and 10:00 h time points in the photoperiod experiment. At 06:00 h, transcript levels for plants under 8L/16D and 24D are expected to match those of 12L/12D, as all were exposed to darkness. Similarly, at 10:00 h, transcript levels for plants under 8L/16D are expected to be same as those under 24D, as these plants were still all exposed to darkness.

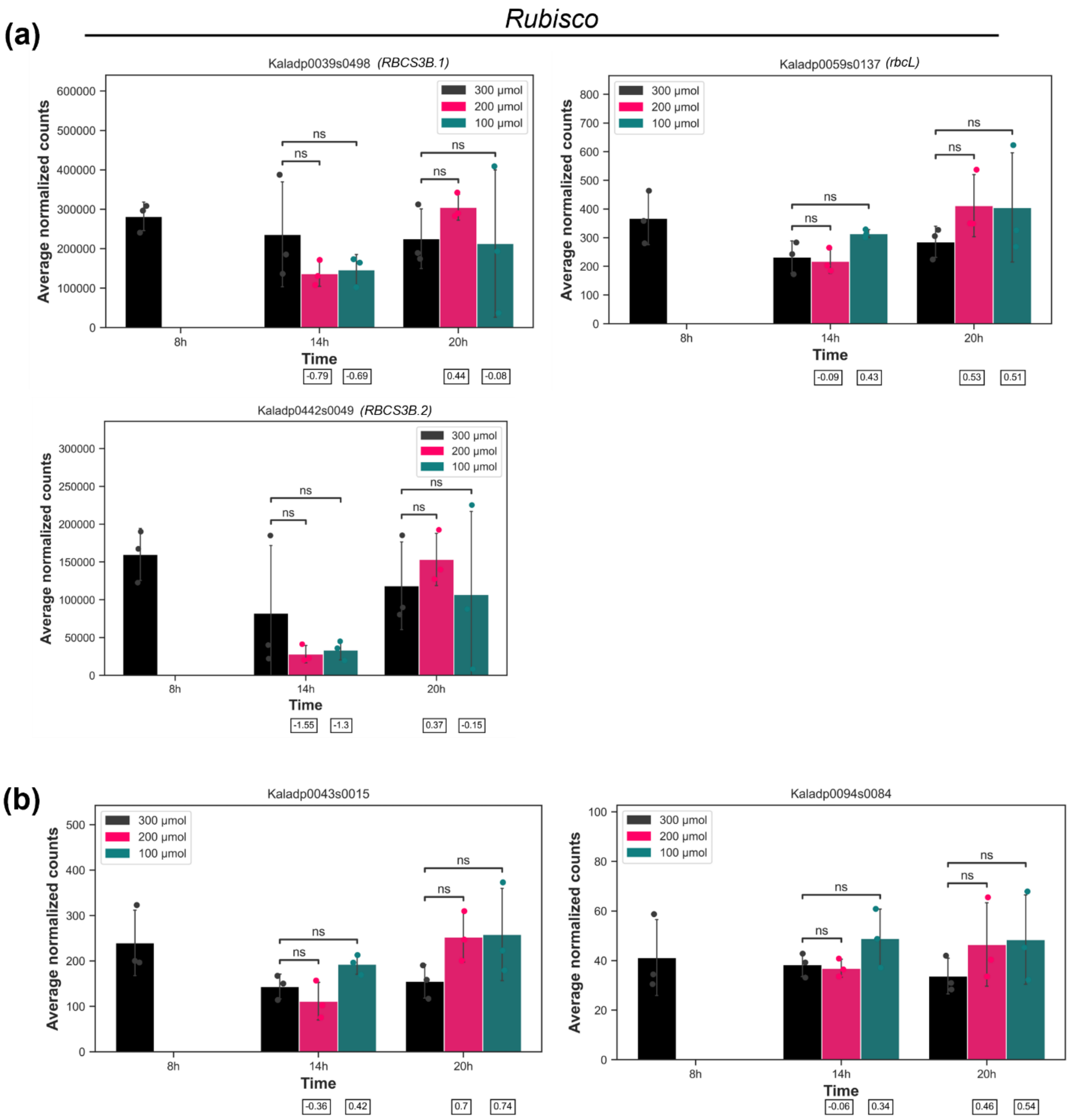
(a)
Rubisco
Kaladp0039s0498 (RBCS3B.1)
Kaladp0059s0137 (rbcL)
Kaladp0442s0049 (RBCS3B.2)
300 µmol
200 µmol
100 µmol
Average normalized counts
Time
8h
14h
20h
ns
-0.79
-0.69
0.44
-0.08
-0.09
0.43
0.53
0.51
-1.55
-1.3
0.37
-0.15
(b)
Kaladp0043s0015
Kaladp0094s0084
-0.36
0.42
0.7
0.74
-0.06
0.34
0.46
0.54

**(c)** *Rubisco*

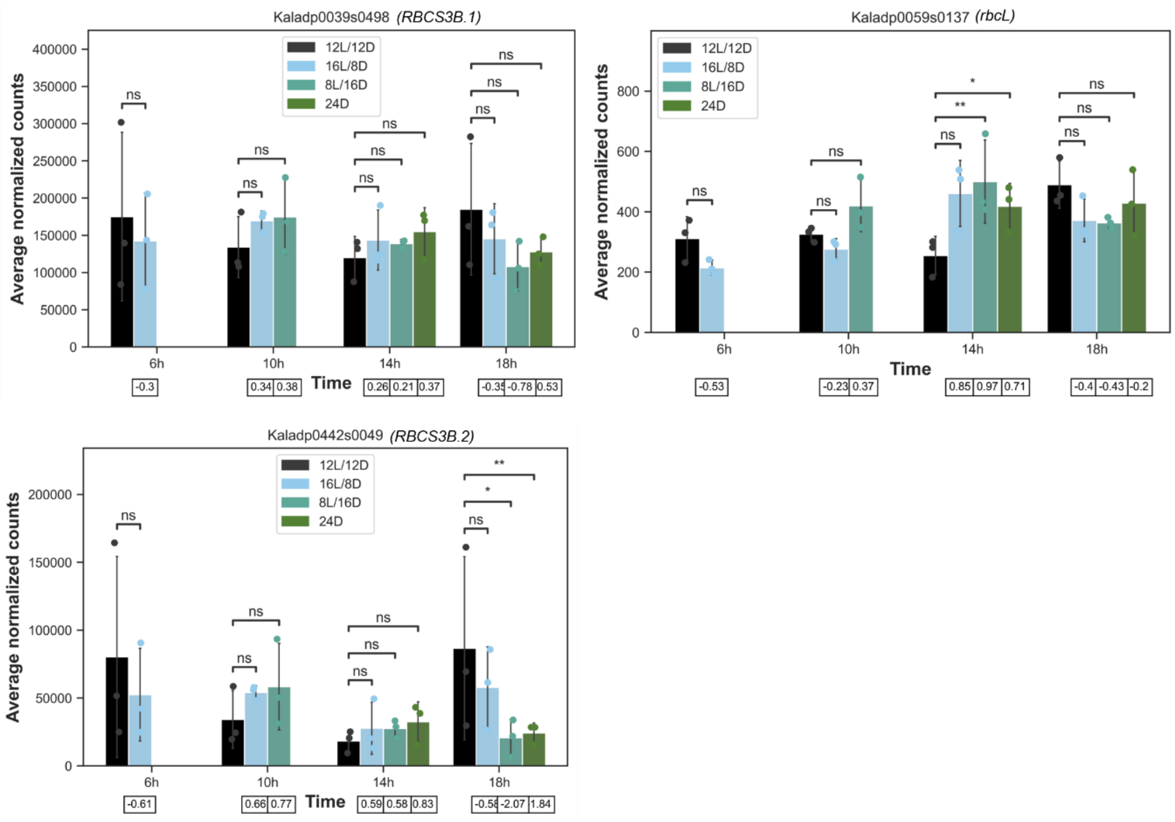

Kaladp0039s0498 (RBCS3B.1)
Kaladp0059s0137 (rbcL)
Kaladp0442s0049 (RBCS3B.2)
12L/12D
16L/8D
8L/16D
24D
Average normalized counts
Time
6h
10h
14h
18h
ns
*
**


**(d)**

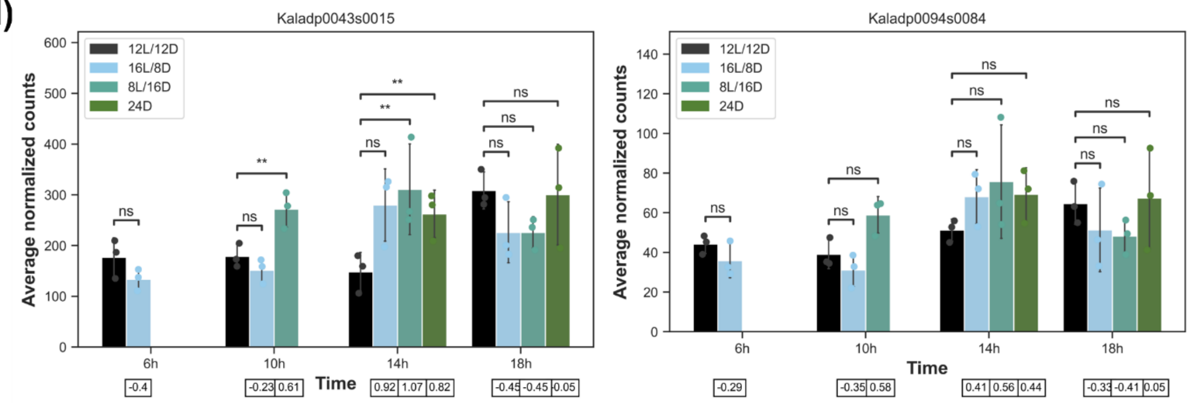

Kaladp0043s0015
Kaladp0094s0084
12L/12D
16L/8D
8L/16D
24D
Average normalized counts
Time
6h
10h
14h
18h
ns
**

**Fig. S5 Temporal transcript abundance (average normalized counts) patterns of *Rubisco activase (RCA)* in leaves of *Kalanchoë fedtschenkoi* under different light intensities and photoperiods.** *RCA* transcript levels for LP5, 6, 7 of *K. fedtschenkoi* under different light intensities (300-200-100 µmol $m^{-2}$ $s^{-1}$) at 08:00, 14:00 and 20:00 h (a) and photoperiods at 06:00, 10:00, 14:00, 18:00 h (b). Data from all *RCA* genes were included in the heatmaps (Fig. 5). Values inside boxes below the bars represent the log2 fold change ($log_2FC$) of the treatment compared to the control (300 µmol $m^{-2}$ $s^{-1}$ in the light intensity experiment and 12L/12D in the photoperiod experiment) (n=3 plants). Error bars represent SD. Significant differences were tested using DESeq2 (padj values) with the significance threshold P=0.05 and are indicated with an asterisk (*P<0.05; **P<0.01; ***P<0.001; ****P<0.0001). RNA-seq samples were collected at 7:45 h in darkness, since lights were switched on at 08:00 h. At this time, plants from the other treatments were also still exposed to darkness and thus received the same treatment. Therefore, data is only available for the control treatment (300 µmol $m^{-2}$ $s^{-1}$) at 08:00 h. The same applies to the 06:00 h and 10:00 h time points in the photoperiod experiment. At 06:00 h, transcript levels for plants under 8L/16D and 24D are expected to match those of 12L/12D, as all were exposed to darkness. Similarly, at 10:00 h, transcript levels for plants under 8L/16D are expected to be same as those under 24D, as these plants were still all exposed to darkness.

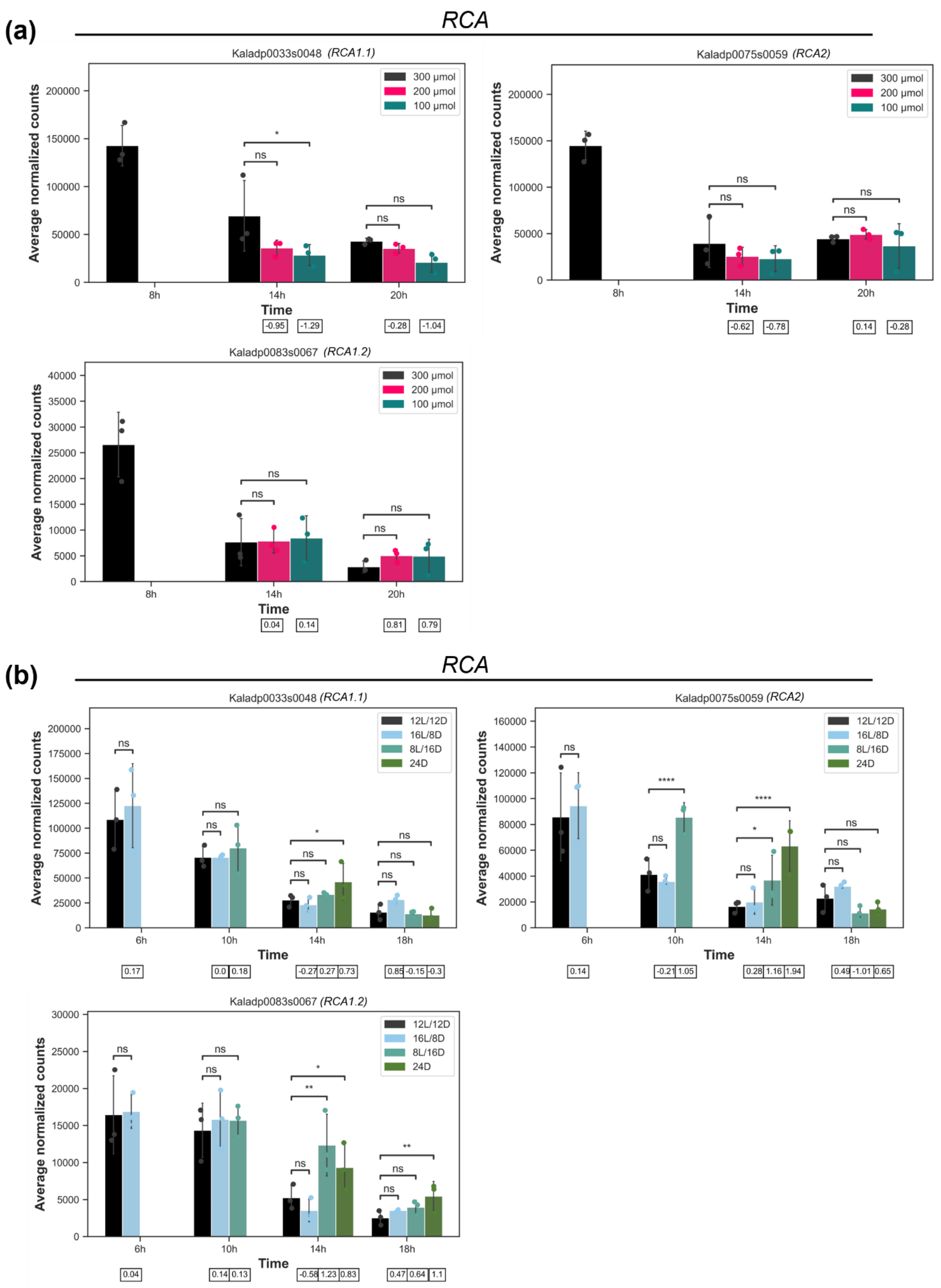
(a)
RCA
Kaladp0033s0048 (RCA1.1)
Kaladp0075s0059 (RCA2)
Kaladp0083s0067 (RCA1.2)
300 μmol
200 μmol
100 μmol
Average normalized counts
Time
8h
14h
20h
-0.95 -1.29 -0.28 -1.04
-0.62 -0.78 0.14 -0.28
0.04 0.14 0.81 0.79
(b)
RCA
Kaladp0033s0048 (RCA1.1)
Kaladp0075s0059 (RCA2)
Kaladp0083s0067 (RCA1.2)
12L/12D
16L/8D
8L/16D
24D
Average normalized counts
Time
6h
10h
14h
18h
0.17 0.0 0.18 -0.27 0.27 0.73 0.85 -0.15 -0.3
0.14 -0.21 1.05 0.28 1.16 1.94 0.49 -1.01 0.65
0.04 0.14 0.13 -0.58 1.23 0.83 0.47 0.64 1.1

**Fig. S6 Temporal transcript abundance (average normalized counts) patterns of *PPDK and PPDK-RP* in leaves of *Kalanchoë fedtschenkoi* under different light intensities and photoperiods.** *PPDK and PPDK-RP* transcript levels for LP5, 6, 7 of *K. fedtschenkoi* under different light intensities (300-200-100 µmol $m^{-2}$ $s^{-1}$) at 08:00, 14:00 and 20:00 h (a, c) and photoperiods at 06:00, 10:00, 14:00, 18:00 h (b, d). Data from both *PPDK* and *PPDK-RP* genes were included in the heatmaps (Fig. 5). Values inside boxes below the bars represent the log2 fold change ($log_2$FC) of the treatment compared to the control (300 µmol $m^{-2}$ $s^{-1}$ in the light intensity experiment and 12L/12D in the photoperiod experiment) (n=3 plants). Error bars represent SD. Significant differences were tested using DESeq2 (padj values) with the significance threshold P=0.05 and are indicated with an asterisk (*P<0.05; **P<0.01; ***P<0.001; ****P<0.0001). RNA-seq samples were collected at 7:45 h in darkness, since lights were switched on at 08:00 h. At this time, plants from the other treatments were also still exposed to darkness and thus received the same treatment. Therefore, data is only available for the control treatment (300 µmol $m^{-2}$ $s^{-1}$) at 08:00 h. The same applies to the 06:00 h and 10:00 h time points in the photoperiod experiment. At 06:00 h, transcript levels for plants under 8L/16D and 24D are expected to match those of 12L/12D, as all were exposed to darkness. Similarly, at 10:00 h, transcript levels for plants under 8L/16D are expected to be same as those under 24D, as these plants were still all exposed to darkness.

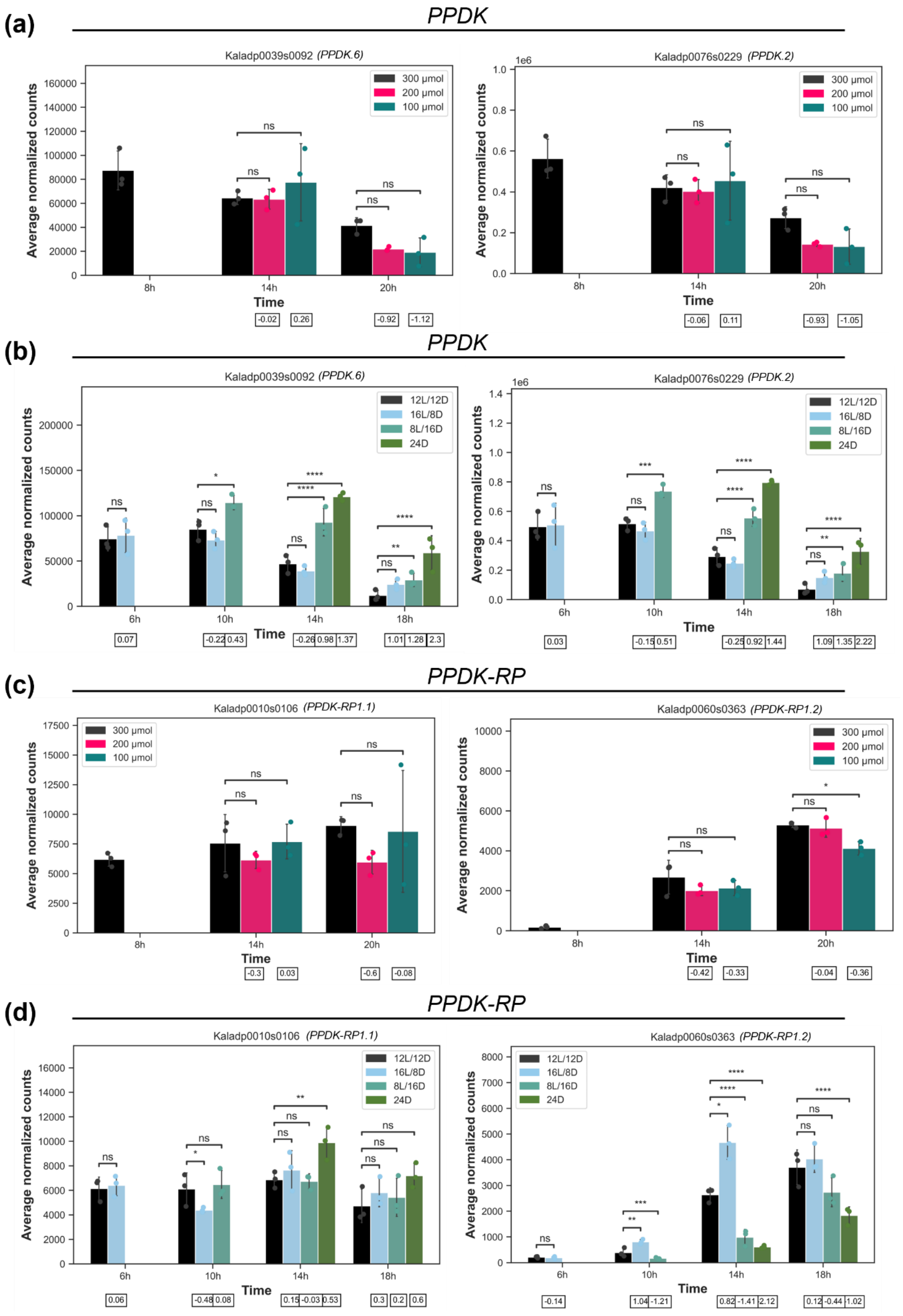

(a)
PPDK
Kaladp0039s0092 (PPDK.6)
Kaladp0076s0229 (PPDK.2)
300 µmol
200 µmol
100 µmol
Average normalized counts
Time
8h
14h
20h
ns
1e6
-0.02
0.26
-0.92
-1.12
-0.06
0.11
-0.93
-1.05
(b)
PPDK
Kaladp0039s0092 (PPDK.6)
Kaladp0076s0229 (PPDK.2)
12L/12D
16L/8D
8L/16D
24D
6h
10h
14h
18h
0.07
-0.22
0.43
-0.26
0.98
1.37
1.01
1.28
2.3
0.03
-0.15
0.51
-0.25
0.92
1.44
1.09
1.35
2.22
(c)
PPDK-RP
Kaladp0010s0106 (PPDK-RP1.1)
Kaladp0060s0363 (PPDK-RP1.2)
-0.3
0.03
-0.6
-0.08
-0.42
-0.33
-0.04
-0.36
(d)
PPDK-RP
Kaladp0010s0106 (PPDK-RP1.1)
Kaladp0060s0363 (PPDK-RP1.2)
0.06
-0.48
0.08
0.15
-0.03
0.53
0.3
0.2
0.6
-0.14
1.04
-1.21
0.82
-1.41
2.12
0.12
-0.44
-1.02

**Table S1 Slope coefficients as a measure for the rates of malate degradation and starch accumulation.** Slope coefficients as a measure for the rates of malate degradation (µmol $g^{-1}$FW $2h^{-1}$) and starch accumulation (µmol Glc eq. $g^{-1}$FW $2h^{-1}$). Linear curves were fitted through metabolite data obtained at time points 10:00h, 12:00h, 14:00h, and 16:00h for leaf pairs 5, 6, or 7 of *Kalanchoë fedtschenkoi* plants exposed to reduced light intensities for one day. A minus sign indicates degradation, while a plus sign denotes accumulation. Data are means ± SE (n=5 plants). Values were compared among the different treatments according to Tukey's HSD test at p<0.05 marked by different letters.

| Treatment | Malate | Starch |
|---|---|---|
| PAR (µmol $m^{-2}$ $s^{-1}$) | Slope/rate | |
| 300 | -10.7 ± 1.7[a] | +10.0 ± 1.3[a] |
| 200 | -8.7 ± 2.7[ab] | +6.0 ± 2.2[b] |
| 100 | -3.2 ± 5.2[bc] | +2.6 ± 1.2[c] |
| 50 | -2.8 ± 3.8[bc] | -0.4 ± 2.3[cd] |
| 10 | -2.1 ± 2.7[c] | -1.0 ± 1.3[d] |

**Methods S1** Extended Materials and Methods.

*Determination of starch content*

Metabolite extraction was conducted by heating frozen, powdered leaf tissue in 80% (v/v) methanol to 80°C for 40 min with intermittent vortexing. The insoluble residue from the methanol extraction was used to determine starch content spectrophotometrically at 340 nm as glucose equivalents (Genesys 10S UV-VIS, Thermo Fisher Scientific, United States), following digestion with a mix of amyloglucosidase (EC 3.2.1.3) and α-amylase (EC 3.2.1.1). The analysis was conducted as earlier described by Ceusters et al. (2008).

*Determination of malic acid content*

Approximately 120 mg of powdered tissue was mixed with 300 µl of ice-cold 4% (v/v) $HClO_4$. The mixture was allowed to thaw slowly on ice for 30 min. The resulting suspension was then centrifuged at 4°C for 10 min at 16 200 *g*. The supernatant from the $HClO_4$ extraction was neutralized at 4°C with 5 M $K_2CO_3$, and the resulting potassium perchlorate precipitate was removed by 10 min centrifugation at 16 200 *g* and 4°C. Five mg activated charcoal was added to the supernatant, and after 15 min at 4°C, removed by new centrifugation. The supernatant was used for measurement of malic acid. Malic acid content was determined using the K-LMAL-116A kit (Megazyme) following the manufacturer's protocol. Malate content was measured in a 500-µl reaction mixture using 6 µL of the initial malic acid extract.

*Determination of sucrose content*

Sucrose was extracted from approximately 100 mg of powdered tissue with 500 µl demineralized water by heating at 60 °C for 30 minutes. After centrifugation at 16 200 *g* for 3 minutes, the resulting sucrose-extract supernatant was stored in a fresh Eppendorf tube and 45 µl of Carrez I and Carrez II was added for the precipitation of proteins, elimination of turbidity and for breaking of emulsions. Discoloration of the extract was initiated by adjusting the pH between 7.5 and 8.5 with 1M NaOH. Subsequently, samples were centrifuged at 16 200 *g* for 20 minutes and the resulting supernatant was used to determine sucrose content via an enzymatic assay kit (K-SUFRG, Megazyme) according to the manufacturer's protocol followed by quantification with the 340 nm light absorbance of NADPH using a spectrophotometer (Genesys 10S UV-Vis, Thermo Fisher Scientific).

*Enzyme activity assays for NAD-ME, NADP-ME, PPDK, and Rubisco*

All extraction steps were performed by homogenizing powdered leaf material at 4 °C in the enzyme specific extraction buffer (approximately 100 mg tissue to 300 µL of extraction buffer). Samples were centrifuged at 16 200 *g* for 2 min at 4 °C in a microfuge. The supernatant was utilized either directly in enzyme assays, or was first desalted by passing twice through a 0.5 ml column of Sephadex-G25 for the NAD-ME assay.

The NAD-ME extraction buffer comprised 100 mM HEPES-KOH, pH 7.0, 2 mM $MnCl_2$, 5 mM DTT, 1% polyvinylpyrrolidone-40, 1 mM EDTA, 2% PEG-20000, and 0.5% Triton X-100. The desalting buffer contained 125 mg/mL Sephadex, 100 mM HEPES-KOH, pH 7.0, 2 mM $MnCl_2$, 5 mM DTT, and 1 mM EDTA. The assay comprised 50 mM HEPES-KOH, pH 7.6, 1 mM EDTA, 1 mM DTT, 5 mM L-malate, 5 mM NAD, 25 µM NADH, 1 U pig heart MDH (Roche Life Sciences), 100 µM acetyl CoA, and 5 mM $MnCl_2$; 25 µM NADH was included to minimize the interference of MDH in the assay and reduce the chance of overestimation of NAD-ME activity (Hatch et al., 1982). After preincubation of extracts for 30 min at 30°C, the reaction was initiated by the addition of 50 µl of extract and change in absorbance at 340 nm was measured for 4 min at 25°C. Preliminary experiments confirmed a linear increase of NADH for at least 6 min.

The NADP-ME extraction buffer comprised 200 mM Tricine-NaOH, pH 7.6, 1 mM EDTA, 2% polyvinylpyrrolidone-40, 2 mM DTT, 1 mM benzamidine hydrochloride, 2% PEG-20000 plus 50 mg $g^{-1}$ tissue $NaHCO_3$, and 200 mg $g^{-1}$ tissue PVPP. The assay comprised 50 mM HEPES-KOH, pH 7.0, 0.1 mM EDTA, 1 mM $NADP^+$, 10 mM L-malate, 5 mM DTT, and 1 mM $MgCl_2$. After preincubation of extracts for 30 min at 30°C, the reaction was initiated by the addition of 100 µl of extract and change in absorbance at 340 nm was measured for 4 min at 25°C. Preliminary experiments confirmed a linear increase of NADPH for at least 6 min.

The PPDK extraction buffer comprised 50 mM HEPES-KOH, pH 8.2, 5 mM DTT, 0.2 mM EDTA, 2% PEG-20000, 2.5 mM $K_2HPO_4$, 2.5 mM pyruvate, and 200 mg $g^{-1}$ tissue PVPP. The assay comprised 50 mM Tris-HCl, pH 8.0, 5 mM DTT, 10 mM MgCl2, 1.25 mM pyruvate, 0.25 mM NADH, 2.5 mM $NaHCO_3$, 2.5 mM $K_2HPO_4$, 1 U pig heart MDH, 1.25 mM ATP, and 6 mM Glc-6-P. The reaction was initiated by the addition of 25 µl of extract and change in absorbance at 340 nm was measured for 4 min at 25°C. Preliminary experiments confirmed a linear decrease of NADH for at least 6 min.

The Rubisco extraction buffer contained 400 mM HEPES-KOH, pH 7.5, 5 mM EGTA, 5 mM $MgCl_2$, 2% (w/v) PEG-20 000, 14 mM β-mercaptoethanol, 16 mg polyvinylpolypyrrolidone (PVPP), and 1 mM PMSF. The initial activity of Rubisco was assayed in a reaction mix that comprised 100 mM Bicine-KOH, pH 8.0, 25 mM $NaHCO_3$, 20 mM $MgCl_2$, 3.5 mM ATP, 3.5 mM P-creatine, 0.25 mM NADH, 5 U 3-phosphoglyceric phosphokinase, 5 U glyceraldehyde 3-phosphate dehydrogenase, and 5 U creatine phosphokinase. After preincubation for 10 min at 25°C, the reaction was initiated by the addition of RuBP to a final concentration of 0.5 mM and change in absorbance at

340 nm was measured for 4 min at 25°C. Preliminary experiments confirmed a linear decrease of NADH for at least 6 min.

*Determination of PPDK and P-PPDK protein abundance via Western blotting*

Total protein extracts of *K. fedtschenkoi* leaves were prepared by mixing approximately 100 mg of $N_2$ crushed tissue with 300 µl of 1x SDS sample buffer [1M Tris-HCl, pH 6.8 at 4 °C, 20% (v/v) glycerol, 4% (v/v) β-mercaptoethanol, and 3% (w/v) sodium dodecyl sulfate (SDS)] in a screw-cap microfuge tube. Samples were boiled immediately for 5 min at 95 °C and were centrifuged at 4 °C for 5 min at full speed. The supernatants were removed and protein contents were determined as described by Bradford (1976) using Bradford reagent (B5702, TCI chemicals).

Proteins were separated on 7.5% polyacrylamide gels (Bio-Rad) and blotted onto 0.2 µm nitrocellulose membranes (Bio-Rad). PPDK protein abundance was determined using antibody purchased from Agrisera (Vännäs, Sweden) (AS13 2647). Phosphorylated PPDK (P-PPDK) antibody [raised against maize (*Zea mays*) PPDK] was kindly provided by Chris J. Chastain, Minnesota State University, Moorhead (Chastain et al., 2018). Goat anti-rabbit IgG HRP conjugate (Agrisera, AS09 602) was used as the secondary antibody. Detection was achieved using the enhanced chemi-luminescence (ECL) system (GE Healthcare) following manufacturer's instructions. Immunoblot signals were detected and digitalized with the Amersham ImageQuant 800 (Cytiva). Duplicate gels were stained with Coomassie Blue to confirm equal loading of the samples and protein integrity.

*Total RNA extraction, RNA sequencing, and bioinformatics*

Total RNA was extracted from 100 mg of liquid nitrogen ground leaf tissue using the Qiagen RNeasy Plant Mini Kit following the manufacturer's protocol with the addition of 13.5 µL of 50 mg/mL polyethylene glycol 20 000 (PEG-20000) to the 450 µL of RLC buffer used for each extraction. Genomic DNA was removed via on-column DNase digestion (Qiagen). RNA quantity, quality, and integrity were assessed by nanodrop (NanoDrop Lite Plus Spectrophotometer, Thermo Scientific), gel electrophoresis, and on the Bioanalyzer (Agilent) respectively. Library preparation and sequencing were performed by Azenta (Genewiz) using the Illumina NovaSeq platform (2x150 bp paired-end sequencing) to obtain 17-55 million reads per sample. Raw sequencing data will be available in a public repository upon acceptance of the article.

RNA-seq data analysis was performed as described in Mohorović et al. (2024), with some modifications for *Kalanchoë fedtschenkoi*. After initial quality check with FastQC, sequence reads were trimmed to remove adapter sequences and bases with poor quality using Trimmomatic v.0.32. The trimmed reads were mapped to the *K. fedtschenkoi* reference genome (Phytozome v.13, https://phytozome-next.jgi.doe.gov/, Yang et al., 2017) using HISAT2. Counting was performed using the HTSeq-count function of HTSeq. A comparison of gene expression between the control and light-treated group of samples was performed in RStudio using DESeq2 (Love et al., 2014). Genes with an adjusted p-value < 0.05 and absolute log2FoldChange > 1 (fold change of 2) were called as differentially expressed genes. Heat-maps and figures were generated in GraphPad Prism 8 and BioRender, respectively. Bar charts with normalized counts were generated using a custom Python script.
